# Multi-model evaluation of phenology prediction for wheat in Australia

**DOI:** 10.1101/2020.06.06.133504

**Authors:** Daniel Wallach, Taru Palosuo, Peter Thorburn, Zvi Hochman, Fety Andrianasolo, Senthold Asseng, Bruno Basso, Samuel Buis, Neil Crout, Benjamin Dumont, Roberto Ferrise, Thomas Gaiser, Sebastian Gayler, Santosh Hiremath, Steven Hoek, Heidi Horan, Gerrit Hoogenboom, Mingxia Huang, Mohamed Jabloun, Per-Erik Jansson, Qi Jing, Eric Justes, Kurt Christian Kersebaum, Marie Launay, Elisabet Lewan, Qunying Luo, Bernardo Maestrini, Marco Moriondo, Gloria Padovan, Jørgen Eivind Olesen, Arne Poyda, Eckart Priesack, Johannes Wilhelmus Maria Pullens, Budong Qian, Niels Schütze, Vakhtang Shelia, Amir Souissi, Xenia Specka, Amit Kumar Srivastava, Tommaso Stella, Thilo Streck, Giacomo Trombi, Evelyn Wallor, Jing Wang, Tobias K.D. Weber, Lutz Weihermüller, Allard de Wit, Thomas Wöhling, Liujun Xiao, Chuang Zhao, Yan Zhu, Sabine J. Seidel

## Abstract

Predicting wheat phenology is important for cultivar selection, for effective crop management and provides a baseline for evaluating the effects of global change. Evaluating how well crop phenology can be predicted is therefore of major interest. Twenty-eight wheat modeling groups participated in this evaluation. Our target population was wheat fields in the major wheat growing regions of Australia under current climatic conditions and with current local management practices. The environments used for calibration and for evaluation were both sampled from this same target population. The calibration and evaluation environments had neither sites nor years in common, so this is a rigorous evaluation of the ability of modeling groups to predict phenology for new sites and weather conditions. Mean absolute error (MAE) for the evaluation environments, averaged over predictions of three phenological stages and over modeling groups, was 9 days, with a range from 6 to 20 days. Predictions using the multi-modeling group mean and median had prediction errors nearly as small as the best modeling group. About two thirds of the modeling groups performed better than a simple but relevant benchmark, which predicts phenology by assuming a constant temperature sum for each development stage. The added complexity of crop models beyond just the effect of temperature was thus justified in most cases. There was substantial variability between modeling groups using the same model structure, which implies that model improvement could be achieved not only by improving model structure, but also by improving parameter values, and in particular by improving calibration techniques.

## 1. Introduction

Crop phenology describes the cycle of biological events during plant growth. These events include, for example, seedling emergence, leaf appearance, flowering, and maturity. Timing of growing seasons and their critical phases as well as estimates of them are increasingly important in changing climate (Olesen et al., 2012, Dalhaus et al., 2018). Matching the phenology of crop varieties to the climate in which they grow is critical for viable crop production strategies (Rezaei et al., 2018, Hunt et al., 2019). Furthermore, accurate simulation of phenology is essential for models which simulate plant growth and yield (Archontoulis et al., 2014; Boote et al., 2010, 2008).

In this study we focus on wheat phenology in Australia. Australia was the world’s ninth largest producer of wheat in 2018 and the sixth largest exporter (Workman, 2020). Crop model predictions of phenology have been used in various studies related to wheat production in Australia. In a study by Luo et al. (2018), the APSIM model was used to simulate changes in phenology, water use efficiency, and yield to be expected from global climate change. The APSIM model was used to evaluate changes in wheat phenology in Australia as a result of warming temperatures in recent decades (Sadras and Monzon, 2006). That model was also used to determine the flowering date at each location associated with highest average yield (Flohr et al., 2017).

Given the interest in using crop models to predict phenology, it is important to evaluate those predictions. How well can wheat phenology be predicted? In trying to answer this question, one must first define exactly what aspect of the models is being evaluated, and then must choose an appropriate methodology for carrying out the evaluation.

It is important to distinguish two different types of model evaluation, which might be termed evaluation of extrapolation predictions and evaluation of interpolation predictions. They differ as to whether or not the data provided for calibration are representative of the target population, i.e. of the range of environments of interest. In one type of study, the objective is to evaluate how well models can extrapolate to conditions not represented in the calibration data. For example, in a multi-model ensemble study on the effect of high temperatures on wheat growth (Asseng et al., 2015), detailed crop measurements were provided for one planting date and the models were evaluated using other planting dates, some with additional artificial heating during growth. The evaluation data thus represented a much larger range of temperatures than represented in the calibration data. This was a test of how well the models can extrapolate to more extreme temperatures than those available for calibration. Other studies have evaluated how well crop models can extrapolate to environments with enhanced CO_2_, given calibration data for current ambient CO_2_ levels (Biernath et al., 2011).

In the second type of study, the calibration data are meant to be representative of the target population. This evaluates how well crop models can generalize from the calibration environments to other similar environments. An example is the study by Ceglar et al. (2019), which used data on wheat phenology under current conditions in Europe for calibration and then predicted phenology for other environments from the same target population. This type of evaluation is adapted, for example, to the case where one has data from a network of variety trials and wants to predict for other sites and years from the same target population, as in Bao et al.. (2017) for yield. It is this aspect of crop phenology models, namely their ability to predict when provided with a sample of data from the target population, that is evaluated in the present study.

A second aspect of evaluation that must be specified is the modeling group or groups that are being evaluated, where modeling group refers to the combination of crop model and the people responsible for running the simulations. We reserve the term “model” specifically for model structure, i.e. the model equations, while modeling group determines both the model structure and the parameter values, which are chosen or estimated by the group running the model. It is clear that predictions depend not only on the model structure but also on the parameter values, so evaluation really refers to the modeling group. Model evaluation studies may refer to a particular modeling group or to an ensemble of modeling groups. Here, we evaluate an ensemble of 28 different modeling groups. The purpose is not to give information about each specific modeling group, but rather to evaluate how well currently active modeling groups can predict phenology for our target population (e.g. what is the error of the best predicting group), how well can one expect a modeling group chosen at random to predict (e.g. what is the mean or median prediction error), and what is the variability between modeling groups (e.g. what is the spread between the best and worst predictors).

It is important to define precisely the evaluation problem (extrapolation or interpolation, single- or multi-group evaluation), but it is also important that the methodology of evaluation be such as to give reliable results. We focus here on the relation of the predictor (model plus parameter values) and evaluation data. It is well-known from statistics that if a predictor is not independent of the evaluation data, then the error for the evaluation data will in general be less than for new environments (Efron, 1986). That is, non-independence in general leads to underestimating prediction errors. The predictor could depend on the evaluation data if, for example, the evaluation data were also used to calibrate the model, or were used to modify the model equations, or were used to tune site characteristics. If the same sites are present in the calibration and evaluation data, then the model has to some extent been tuned to those sites, and so the predictor is not independent of the evaluation data even if the evaluation data have not been used directly to fit the model. Having the same sites in the calibration and evaluation data is often the case for evaluation studies (Andarzian et al., 2015; Asseng et al., 2008; Chauhan et al., 2019; Hussain et al., 2018; Yuan et al., 2017).

There do not seem to have been any evaluation studies of prediction of wheat phenology in Australia based on results from multiple modeling groups, where the calibration data are sampled from the target population (i.e. evaluation of interpolation predictions). The purpose of this study is to present such an evaluation, using a rigorous approach where the parameterized model is independent of the evaluation data.

## 2. Materials and Methods

### 2.1 Experimental data

The data are a subset from a multi-cultivar, multi-location, and multi-sowing date trial for wheat in Australia, described in Lawes et al. (2016). The environments reflect the diversity in the wheat-growing regions of Australia (Fig. 1). Only the data for cultivar Janz, classified as a fast-moderate maturing cultivar, were used here. The data are from 10 sites, located throughout the grain growing region each with one to three sowing years and three planting dates in each year (overall 66 environments, i.e. site-sowing date combinations, Table 1). The sowing dates at each site correspond to early, conventional, and late sowing. Plant density was 100-120 plants/m^2^, and sowing depth was 20-35 mm. Nutrients were managed to be non-limiting. There were 1-3 repetitions for each environment (average of 2.1 repetitions).

**Figure 1.**
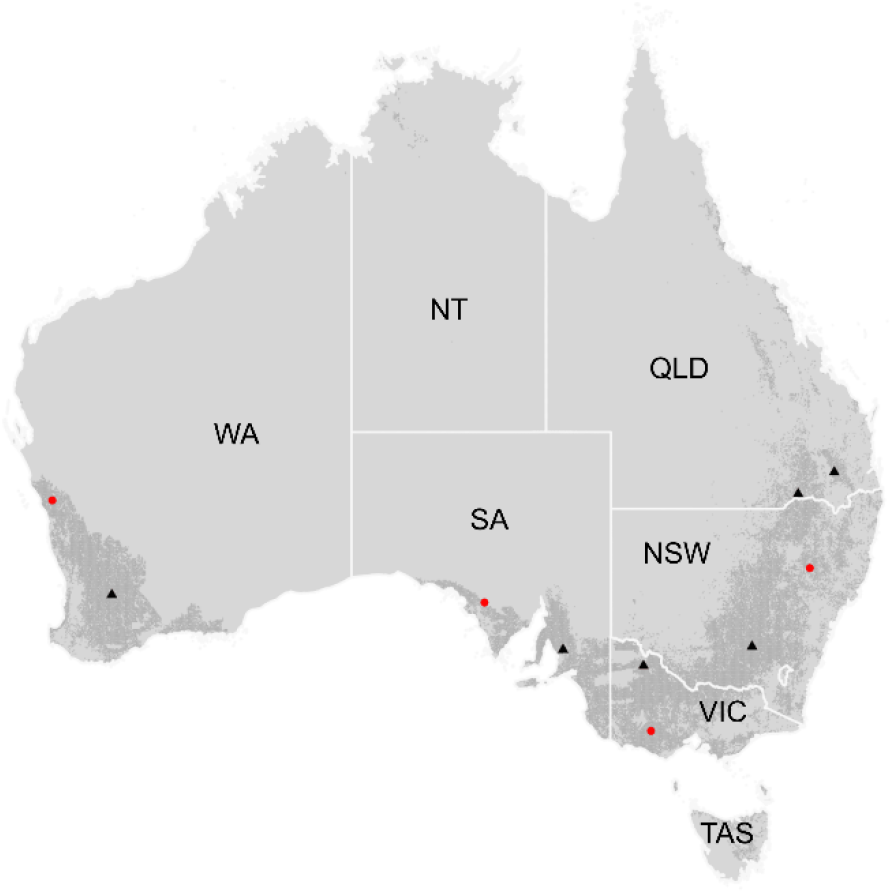
Location of calibration (red circles) and evaluation (blue triangles) sites across the Australian cropping zones (shaded area; Source: Teluguntla et al., 2018).

**Table 1.** Sites and sowing dates for calibration (underlined) and evaluation (bold). Note that the calibration and evaluation data have neither sites nor years in common.

| site\ year | 2010 | 2011 | 2012 |
| --- | --- | --- | --- |
| Bungunya |  |  | **2012-05-10** |
| (Queensland) |  |  | **2012-05-22** |
|  |  |  | **2012-06-23** |
| Corrigin<br>(West Australia) |  |  | **2012-05-02**<br>**2012-05-21**<br>**2012-06-21** |
| Eradu<br>(West Australia) | <u>2010-05-14</u><br><u>2010-05-27</u><br><u>2010-06-22</u> | <u>2011-04-29</u><br><u>2011-05-24</u><br><u>2011-06-23</u> |  |
| LakeBolac<br>(Victoria) | <u>2010-05-03</u><br><u>2010-05-19</u><br><u>2010-07-08</u> | <u>2011-05-09</u><br><u>2011-06-03</u><br><u>2011-06-16</u> |  |
| Minnipa<br>(South Australia) | <u>2010-04-30</u><br><u>2010-05-31</u><br><u>2010-06-24</u> | <u>2011-05-13</u><br><u>2011-05-27</u><br><u>2011-06-24</u> |  |
| Nangwee<br>(Queensland) |  |  | **2012-05-17**<br>**2012-05-31**<br>**2012-06-23** |
| Spring Ridge<br>(New South Wales) | <u>2010-05-10</u><br><u>2010-06-11</u><br><u>2010-07-01</u> | <u>2011-05-09</u><br><u>2011-06-06</u><br><u>2011-06-23</u> |  |
| Temora<br>(New South Wales) |  |  | **2012-05-05**<br>**2012-05-23**<br>**2012-06-25** |
| Turretfield<br>(South Australia) |  |  | **2012-05-30**<br>**2012-06-15**<br>**2012-07-05** |
| Walpeup |  |  | **2012-04-27** |
| (Victoria) |  |  | **2012-06-04** |
|  |  |  | **2012-07-18** |

Plots were visited regularly (about every two weeks) starting soon after emergence of the early sowing and ending after crop maturity, and the Zadoks growth stage (Zadoks et al., 1974), on a scale from 1-100, was determined. Overall, there were 709 combinations of environment and measurement date, with an average of 10.7 stage notations per environment. The stages to be predicted here are stage Z30 (Zadoks stage 30, pseudostem, i.e. youngest leaf sheath erection), stage Z65 (Zadoks stage 65, anthesis half-way, i.e. anthers occurring half way to tip and base of ear), and stage Z90 (Zadoks stage 90, grain hard, difficult to divide). These stages are often used for management decisions or to characterize phenology.

In preparing the data for the simulation study, a linear interpolation was performed between each pair of stages, to give the date for every integer Zadoks stage from the first to the last observed stage. At 10 of the 709 measurement dates, observed Zadoks stage decreased slightly (by an average of 3 on the Zadoks scale) compared to the previous date, due to sampling variability. In that case both observed Zadoks stages were replaced by the average for the two dates, before interpolation. The interpolated values were provided in order to avoid different modeling groups using different methods for interpolating the data, which would have added additional uncertainty unrelated to the model performance.

The average standard deviation of observed Zadoks stages based on the replicates was 0.93 days. The standard deviation of interpolated days after sowing to Z30, Z65, and Z90 was calculated using a bootstrap. For a day with *r* replicates, a sample of size *r* was obtained by drawing values at random with replacement, independently for each measurement date. Then the Zadoks values were interpolated as for the original data. This was done 1000 times, giving standard deviations of 1.8 days for observed days to Z30, 0.9 days for observed days to Z65, and 0.5 days for observed days to Z90, respectively.

Part of the data was provided to the modeling groups for calibration, and part was never revealed to participants and used for evaluation. The calibration data originated from four sites, two years, and three planting dates, so overall 24 environments. The evaluation data were from six sites, one year, and three planting dates for a total of 18 environments (Table 1). Dates of Z30, Z65 and Z90 were observed at respectively 16, 18 and 5 of these 18 environments. The data were divided in such a way that the calibration and evaluation data had neither sites nor years in common.

To characterize the environments, we calculated for each environment the average temperature from sowing to Z30, Z65, and Z90, the average photoperiod from Z30 to Z65 using the daylength function in the R package insol (Corripio, 2019.; R Core Team, 2017) and days to full vernalization using the model in van Bussel et al. (2015) with a required duration of exposure to vernalizing temperatures (*V_sat_*) of 25 days, estimated from the figure in their paper. Figure 2 shows the range of average temperature, day length, and days to vernalization for the calibration and evaluation environments as well as the range of observed calendar days to Z30, Z65, and Z90. The range of values for the evaluation data is always within the range of the calibration data, with the single exception of photoperiod. While the median and maximum day lengths were very similar for the two sets of environments, the shortest day length was 11.5 hours among calibration environments, while among the evaluation environments the shortest day length was 10.1 hours.

**Figure 2.**
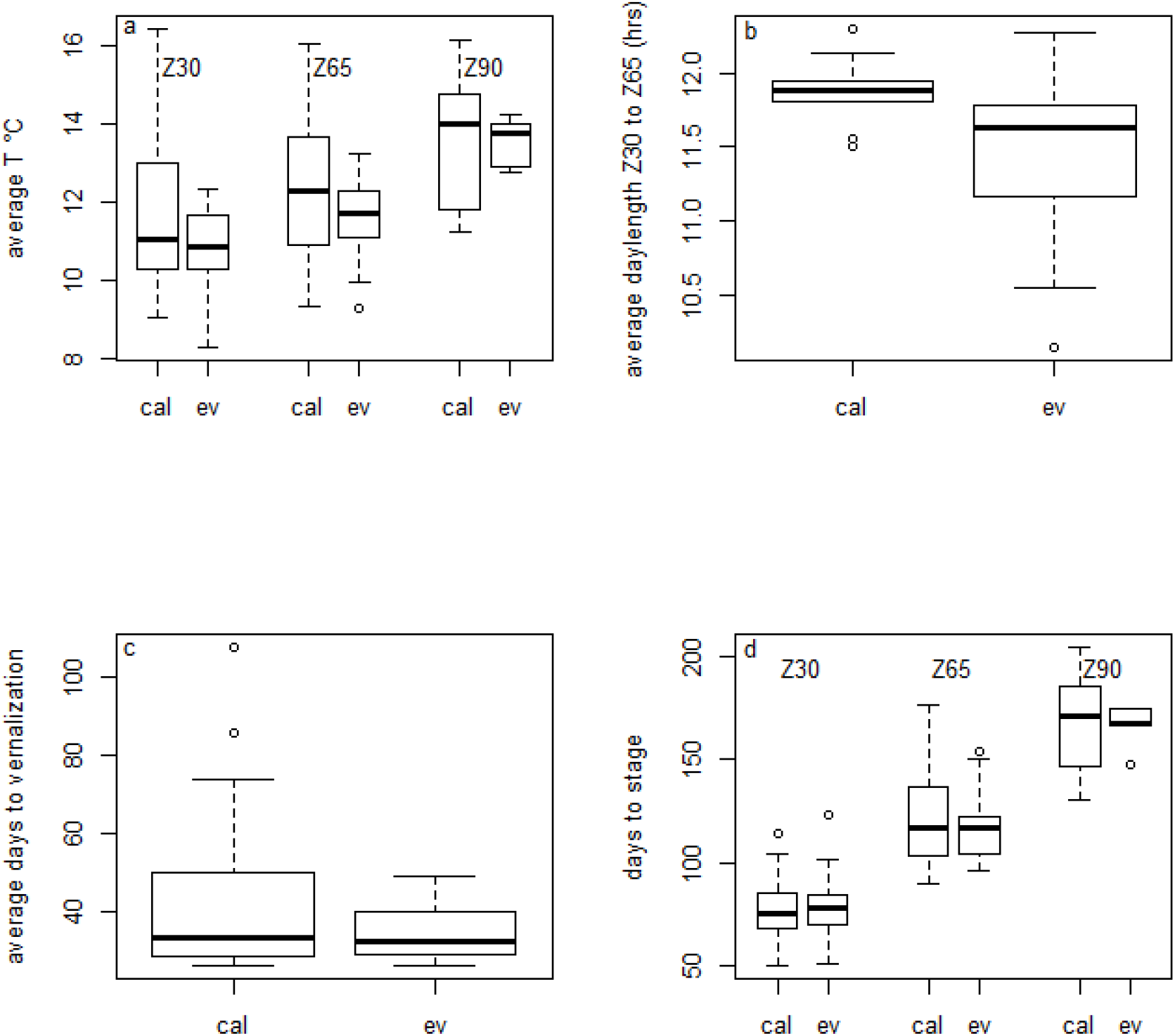
Boxplots of a) average temperatures from sowing to Zadoks stages Z30, Z65, and Z90 b) average day length between observed days of Zadoks stages Z30 and Z65 c) average days from sowing to complete vernalization d) average days from sowing to Zadoks stages Z30, Z65, and Z90. Results are shown separately for the calibration (ca) and evaluation (ev) environments. Boxes indicate the lower and upper quartiles. The solid line within the box is the median. Whiskers indicate the most extreme data point which is no more than 1.5 times the interquartile range from the box, and the outlier dots are those observations that are beyond that range.

### 2.2 Modeling groups

Twenty-eight different modeling groups participated in this study, where modeling group refers to the group of people conducting the modeling exercise. Each modeling group is associated with some specific model structure (some specific named model) and also with some specific parameter values. The model structures involved are presented in Supplementary Table S1. Models were considered to have the same structure even if the version number was different, because version differences are expected to be negligible for phenology. Three of the model structures were used by more than one group. Since different groups using the same structure obtained different results, identifying the contributions by the name of the model would be misleading. Furthermore, the performance of specific groups was not of major interest here. Therefore the modeling groups were anonymized, and only identified by a number. There is no model M5 because that group dropped out in the course of the study. The model structures used by more than one group are noted S1 (three groups), S2 (three groups) and S3 (two groups).

Details about the way phenology is modeled by each model structure can be found in the references for each model (Supplementary Table S1). Here we give only a brief overview. The principal factors that affect winter wheat developmental rate are temperature, day length and degree of vernalization (Johnen et al., 2012). Most, but not all, model structures take into account all three factors. The simplest approach to modeling the effect of temperature is to assume that development rate increases linearly with daily average temperature above some base temperature (a parameter). In other models the rate may be constant above some optimal temperature (a parameter), development rate may decline above the optimum temperature at some rate (a parameter), or development rate may be some more complex function of temperature (Kumudini et al., 2014; Wang et al., 2017). The parameters of the temperature response curve may differ depending on development stage. The effect of photoperiod on development rate is often modeled as a multiplier that is a piecewise linear function of photoperiod. The function increases with some slope (a parameter) up to a threshold photoperiod (a parameter), and then is 1 for photoperiods longer than the threshold. Vernalization, which must be accomplished before the plant can flower, requires a period of cold temperatures. Vernalization parameters can include the upper limit for temperature to count as vernalizing, and the required number of vernalizing days. Some models also relate development to the rate of leaf appearance (called the phyllochron, a parameter) or rate of tillering. Finally, several models also take into account the effect of cold or drought stress on development rate. If drought stress is taken into account, then development rate is related to all the processes that determine soil moisture and plant water uptake.

The multi-model ensemble here was an “ensemble of opportunity” meaning that any modeling group that asked to join was accepted. The activity was announced on the list server of the Agricultural Modeling Inter-comparison and Improvement Project (AgMIP) and on the list servers of several models. In addition to the original models, we defined two ensemble models. The model e-mean has predictions equal to the mean of the simulated values. The model e-median has predictions equal to the median of the simulated values.

### 2.3 Simulation experiment

Each participating modeling group was provided with weather, soil, and management data for all environments, as well as all available observed and interpolated values for days to each Zadoks stage for the calibration data. Participants were requested to return simulated values for number of days from sowing to emergence (even though days to emergence was never observed) and values for number of days from sowing to stages Z30, Z65, and Z90 for all environments, including both the calibration environments and the evaluation environments.

### 2.4 Evaluation

As our basic metric of model error, we use the mean absolute error (MAE). For a model *m*, MAE is

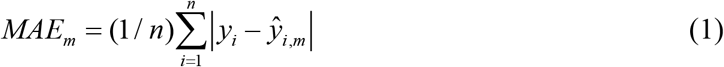

where *y_i_* is the observed value for environment *i* and 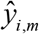 is the value simulated by modeling group *m* for that environment. The sum is over either calibration environments, to evaluate goodness-of-fit, or over evaluation environments, to estimate prediction error. This is preferred over mean squared error (MSE) or root mean squared error (RMSE), because unlike MSE, MAE does not give extra weight to large errors (Willmott and Matsuura, 2005). To test whether MAE is the same for prediction of days to different stages, we used the R function pairwise.t.test, with method=”holm” to correct for multiple comparisons. We also calculated MSE, RMSE, and NRMSE (normalized root mean squared error) for comparison with other studies.

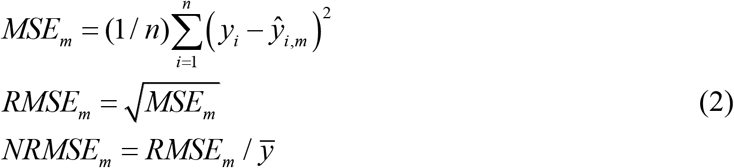

where 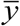 is the average of the observed values.

We considered two skill measures. A skill measure compares prediction error of the modeling group to be evaluated with the error of a simple model used for comparison. We define two simple models, and therefore two skill measures. Both use MSE, rather than MAE, as the measure of model error, in keeping with usual practice. The first simple model, noted “naive”, predicts that days to each stage will be equal to the average number of days to that stage in the calibration data. The predictions of the naïve model here are 77.1, 123.1, and 166.5 days from sowing to stages Z30, Z65, and Z90, respectively. The first skill measure, modeling efficiency (EF), is defined as

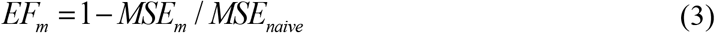

The naive model ignores all variability and predicts that days to any stage will be the same regardless of the environment. A model with EF ≤ 0 is a model that does no better than the naive model, and so would be considered a very poor predictor. A perfect model, with no error, has modeling efficiency of 1. Often modeling efficiency is based on the fit of a calibrated model to the data used for calibration (McCuen et al., 2006). Here, in contrast, the naïve model is based on calibration data and used to predict for independent data.

The naïve model is a very low baseline for evaluating a crop model. We therefore introduce a more realistic, but still simple model which takes into account the effect of temperature on phenology. This “onlyT” model predicts that degree days (°D) from sowing to each stage will be equal to the number of degree days from sowing to that stage in the calibration data, where degree days on any calendar day is equal to average temperature that day. The predictions of the onlyT model are that Z30 will occur 893.7 °D after sowing, Z65 will occur 1476.0 °D after sowing, and Z90 will occur 2245.7 °D after sowing. The second skill measure, noted skillT, is then

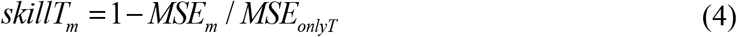

where *MSE_onlyT_* is MSE for the onlyT model. As for any skill measure, a perfect model has skillT = 1 and a model that does no better than the onlyT model has skillT ≤ 0

### 2.5 Sources of variability

A major interest of ensemble studies is that they provide information on the variability in simulation results between different modeling groups. This variability can arise from differences in model structure between different modeling groups or differences in parameter values for groups that use the same model structure. In this study, three of the model structures are used by more than one modeling group. This makes it possible to estimate separately the variance in simulated values due to structure and the variance due to modeling group nested within structure (i.e. due to differences in parameter values). We treat the simulated values as a sample from the distribution of plausible model structures and plausible parameter values. According to the law of total variance (Casella and Berger, 1990), the total variance of simulated values can be decomposed into two parts as

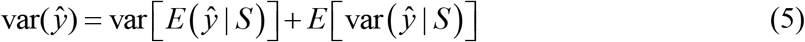

where 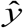 are the simulated values, *S* is model structure, E is the expectation, var is the variance, and the notation |S means that the expectation (in the first term on the right hand side) or the variance (in the second term on the right hand side) is taken separately for each value of model structure. We estimated the first term by first calculating the average simulated value for each structure (if a structure is represented by a single modeling group, this is just the value simulated by that group), and then calculating the variance of those average values. This is the between-structure variability. To estimate the second term, we first calculated the variance between simulated values for each of the three structures with multiple groups. Then we calculated the average of those variances. This is the within-structure variability (i.e. variability due to parameters).

## 3. Results

### 3.1 Prediction error and skill

MAE values for the evaluation data are shown in Figure 3 and summarized in Table 2. Results for individual modeling groups are given in Supplementary Table S2. Median MAE values (and ranges) were 12 days (8-25 days) for days to Z30, 10 days (5-24 days) for days to Z65, and 7 days (1-22 days) for days to Z90. The median (and range) of MAE averaged over the three stages was 9 days (6-20 days). The ensemble predictors e-mean and e-median both had averaged MAE values of 7 days. They were both only marginally worse than the best two individual modeling groups, and e-median was marginally better than e-mean. For comparison with other studies, we also report other criteria of error in Table 2.

**Figure 3.**
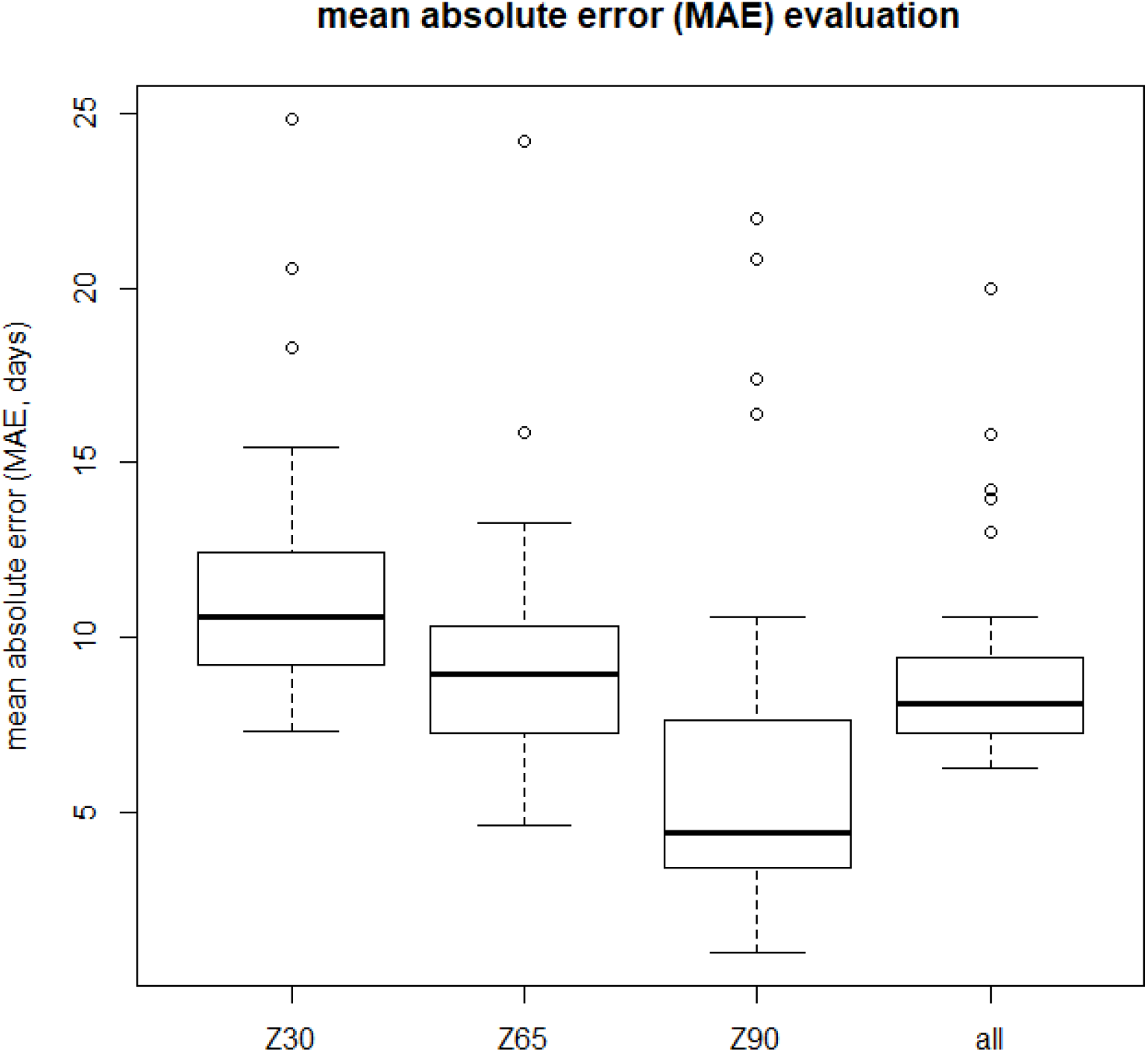
Boxplot of mean absolute error (days) for each development stage and averaged over stages, for the evaluation data. The variability is between different modeling groups. Boxes indicate the lower and upper quartiles. The solid line within the box is the median. Whiskers indicate the most extreme data point which is no more than 1.5 times the interquartile range from the box, and the outlier dots are those observations that are beyond that range.

**Table 2.**
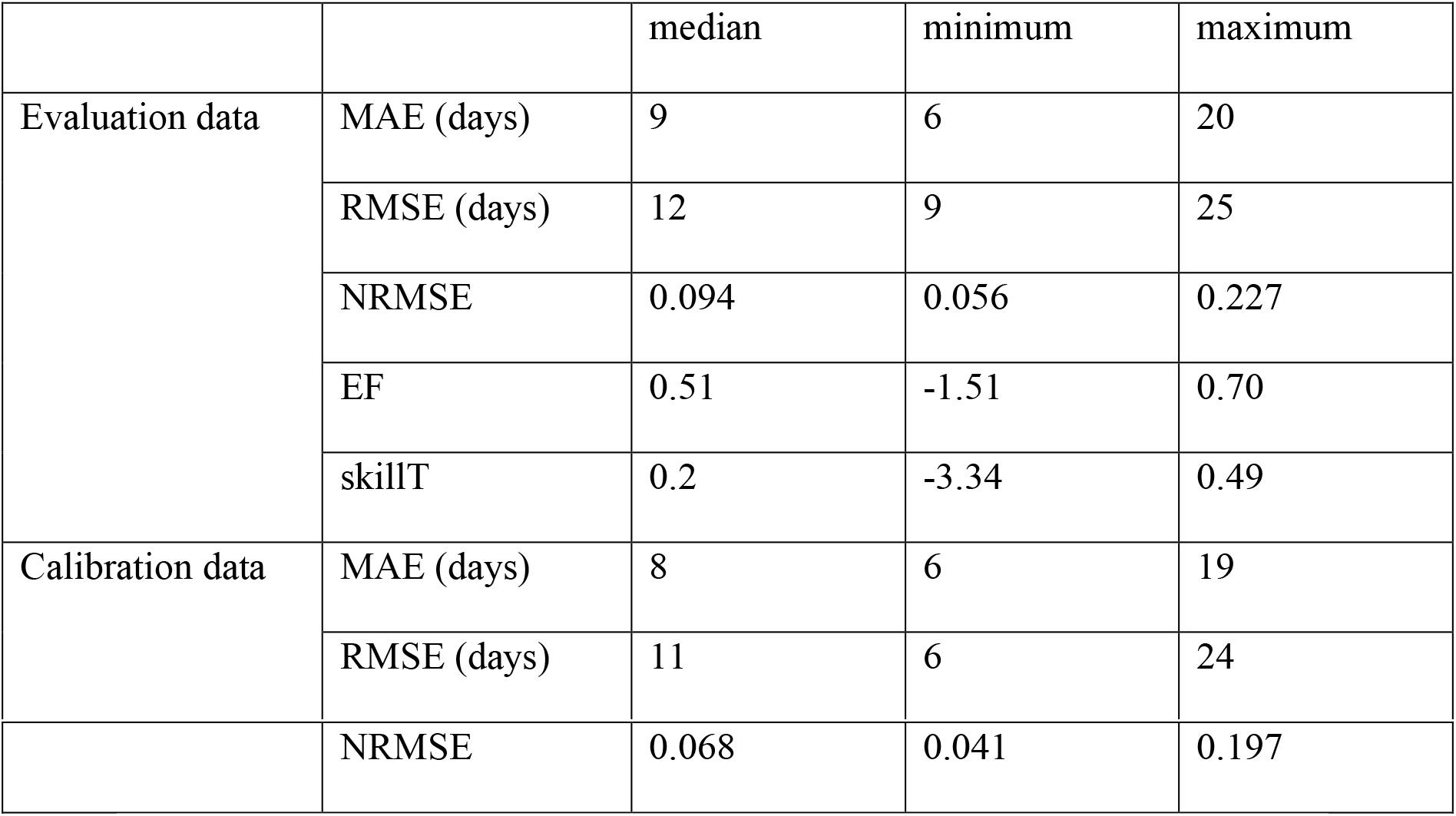
Summary of prediction errors for the evaluation and calibration environments, in each case averaged over predictions of days to stages Z30, Z65, and Z90 except for NRMSE, where the values refer to predictions of number of days to stage Z65. The median, minimum, and maximum error over modeling groups are shown.

|  |  | median | minimum | maximum |
| --- | --- | --- | --- | --- |
| Evaluation data | MAE (days) | 9 | 6 | 20 |
|  | RMSE (days) | 12 | 9 | 25 |
|  | NRMSE | 0.094 | 0.056 | 0.227 |
|  | EF | 0.51 | -1.51 | 0.70 |
|  | skillT | 0.2 | -3.34 | 0.49 |
| Calibration data | MAE (days) | 8 | 6 | 19 |
|  | RMSE (days) | 11 | 6 | 24 |
|  | NRMSE | 0.068 | 0.041 | 0.197 |

Boxplots of EF and skillT for the evaluation data are shown in Figure 4. The median EF value of the individual modeling groups, averaged over stages, was 0.51, and 86 % of the modeling groups had EF > 0. The median skillT value of the individual modeling groups, averaged over stages, was 0.20, and 68% of the modeling groups had skillT > 0.

**Figure 4.**
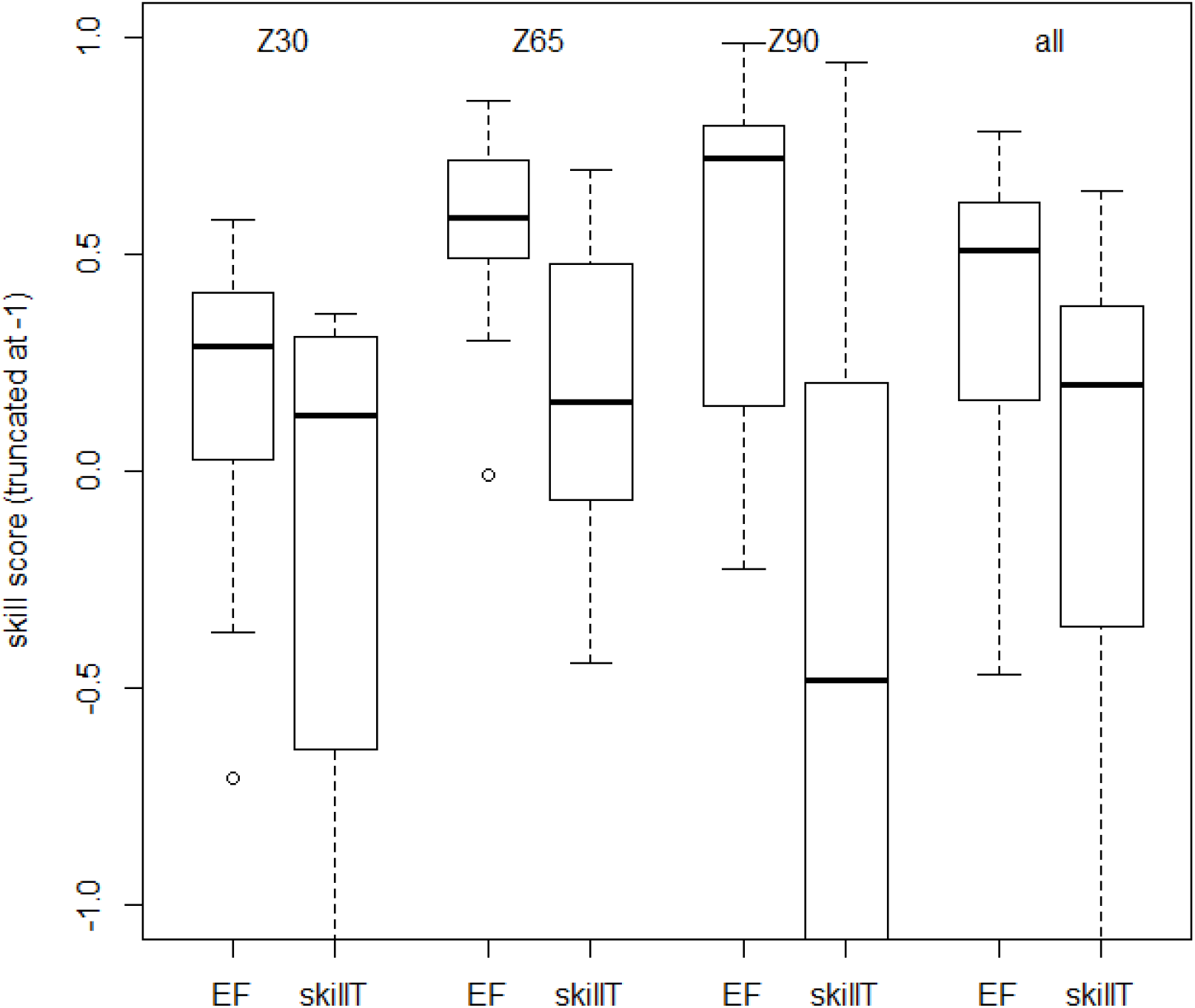
Boxplots of skill scores for prediction of days to Zadoks stages Z30, Z65, and Z90, and averaged over stages (all) for the evaluation data. Skill score is 1 for a modeling group that predicts perfectly, and is less than or equal to 0 for a modeling group that does no better than using average days to each stage in the calibration data (EF skill score) or than using the average number of degree days to each stage in the calibration data (skillT skill score). Boxes indicate the lower and upper quartiles. The solid line within the box is the median. Whiskers indicate the most extreme data point which is no more than 1.5 times the interquartile range from the box, and the outlier dots are those observations that are beyond that range. For readability the y axis is cut off at −1.

Overall MAE for the evaluation data and the calibration data for the same modeling group were correlated. The calibration value explains 46 % of the variability in the evaluation data (R^2^ = 0.46).

### 3.2 Sources of variability

There was substantial variability between modeling groups for each individual prediction, including between modeling groups that share the same model structure (Supplementary Figure S1). Averaged over the evaluation environments and over all three stages Z30, Z65, and Z90, the estimated within-structure standard deviation was 4.3 days and the estimated between-structure standard deviation was 11.9 days, so the within-structure standard deviation was 36 % as large as the between-structure standard deviation.

## 4. Discussion

### 4.1 Comparison of calibration and evaluation environments

The calibration and evaluation environments were drawn from the same target population, namely wheat crops in the major wheat growing regions in Australia, with current climate and local management practices. We compared the calibration and evaluation environments for the main characteristics that are likely to affect phenology, namely temperature, day length, and accumulation of vernalizing temperatures. Temperatures and vernalizing durations of the evaluation environments were within the ranges of the calibration environments, but the evaluation data had a larger range of day lengths than the calibration data. This is the result of sampling variability, and may have led to larger prediction errors than if the calibration data had a range of day lengths comparable to that of the evaluation data. However, the range of days to each phenology stage for the evaluation data was always within the range for the calibration data. We conclude that this study represents a case where the calibration and evaluation data represent a similar range of conditions (with the caveat just mentioned concerning photoperiod). This type of situation is of particular importance, for example, where one wants to calibrate a crop model using current conditions and subsequently test possible sowing dates within a limited range, or to compare phenology of multiple potential cultivars at specific sites within the calibration domain.

### 4.2 Prediction error

The evaluation here was based on data which had neither sites nor years in common with the calibration data. This was thus a rigorous estimate of how well crop modeling groups can predict wheat phenology for unseen sites and weather, when provided with calibration data sampled from the target population. The median MAE among models averaged over phenology stages was 9 days, which was substantially larger than the standard deviation of observed stages, which was in the range 1-2 days. The best modeling group had an average MAE of 7 days, which was still substantially larger than the standard deviation of observed stages. MAE values were significantly larger for prediction of days to Z30 than for prediction of days to later Zadoks stages. This may be due to the large variability between groups in predicting time to emergence, which is discussed in more detail below. Time to emergence is a major part of the time to Z30, but a smaller fraction of time to Z65 or Z90.

Chauhan et al. (2019) reported a value of NRMSE of 0.062 for prediction of time to flowering of wheat in Australia, for a version of APSIM taking the effect of water stress on phenology into account. In that study, the model was adjusted to some extent to the data used for evaluation, so the reported error probably underestimates the error for new environments. That reported value was in any case within the range of NRMSE values found for different modeling groups here, for both the evaluation data (NRMSE here from 0.056 to 0.227) and the calibration data (NRMSE here from 0.041 to 0.197). Asseng et al. (2008), using the APSIM model, found RMSE of 4 days for wheat phenology predictions (mostly predictions of days to anthesis) for 44 different environments in Western Australia, a level of error which was smaller than the minimum RMSE of 9 days found here for the evaluation data, and even smaller than the minimum RMSE of 6 days found here for the calibration data. In that study, the phenology model was again adjusted to some extent to the data (S. Asseng, 2020, pers. comm.), which could explain the smaller errors.

The above comparisons suggest that prediction errors are very roughly similar between studies, but that there are differences depending on the details of the prediction problem and the way prediction error is evaluated. It is clearly useful to build up a knowledge base concerning phenology prediction error, as a baseline for comparison for future studies or even as a default value if evaluation is not done. Contributions to the knowledge base will be all the more useful, to the extent that the details of the prediction problem are clearly specified (including whether it is of type interpolation or extrapolation and including a characterization of the target population) and to the extent that the evaluation has a rigorous separation between the predictor and the evaluation data. The present study should therefore be a valuable contribution to such a knowledge base.

It is of interest to compare the results here with those from a study structured like the present study (calibration and evaluation environments with similar characteristics, evaluation data not used for model development or tuning) but where the evaluation concerned prediction of two phenological stages of wheat in France, namely BBCH30 (equivalent to Z30) and BBCH55 (equivalent to Z55) (Wallach et al., 2019). To a large extent, the same modeling groups participated in both studies. Specifically, the French study included 27 different modeling groups, 26 of which participated in the present study. A comparison between the two studies gives an indication of variability in prediction error for the same modeling groups but for different target populations (Australian wheat in one case, French wheat in the other) and for somewhat different calibration data and predicted stages.

MAE averaged over the evaluation environments and over predicted stages ranged from 3 to 13 days (median 6 days) for the French data compared to 6 to 20 days (median 9 days) for the Australian data. The target population (wheat fields in Australia versus wheat fields in France) thus had a substantial effect on prediction errors. A detailed analysis of the underlying reasons for the larger errors in Australia is beyond the scope of this study. However, one possible contributing cause is the simulation of time to emergence. The average simulated time to emergence for all French environments was 10 days after sowing, and the mean standard deviation between modeling groups was 4 days. The corresponding values for the Australian environments were a mean emergence time of 15 days after sowing, and a mean standard deviation between modeling groups of 18 days. This very large standard deviation for the Australian environments, pointing at major differences between modeling groups, may be due to dry conditions in some environments and the uncertainty regarding initial soil conditions, leading some models to simulate very long times to emergence (up to 107 days, Supplementary Figure S1). This suggests that for Australian environments, it would be valuable to have observations of time to emergence for calibration. It seems that for many modeling groups, it would be worthwhile to revisit the predictions of time to emergence under conditions like those of the Australian environments, taking advantage of specific modeling studies of time to emergence for wheat (Lindstrom et al., 1976; Wang et al., 2009).

An important question in modeling is whether the same modeling groups perform best for all target populations, or whether different groups are best for different target populations. There is quite a bit of scatter in the graph of MAE for the Australian versus French environments (Supplementary Fig. S2), but the rank correlation between the two (Kendall’s tau) is 0.31, which is statistically significant (*p*=0.013). This suggests that there are modeling groups which perform better than others over a wide range of environments. Once again, it is prudent to repeat that this applies to the case where calibration is based on environments that are sampled from the target distribution. Prediction errors for extrapolation to conditions very different than those of the calibration data might behave very differently.

### 4.3 Skill measures

While prediction error is of course of interest, skill scores may be even more useful, as they indicate how models compare to alternative methods of prediction. Note that the EF skill score used here is somewhat different than the usual definition. Here, the naïve model is based solely on the calibration data, so this is in fact a feasible predictor. The more usual definition of the naïve model is the mean of all the data, including the data used for evaluation. Overall, all except four modeling groups had smaller MSE (were better predictors) than the naïve model.

The EF criterion is a rather low baseline for evaluating the usefulness of crop models for predicting phenology. Our second skill measure compares model MSE and MSE of the onlyT model, which assumes a constant number of degree days from sowing to each Zadoks stage, and estimates that number based on the calibration data. This should be a better predictor than the naïve model if photoperiod and vernalization effects are limited, and so is a more stringent test of usefulness of process models. We found that the onlyT model was indeed a better predictor than the naïve model. Nonetheless, 19 of the modeling groups performed better than the onlyT model. It seems that in most cases here, the added complexity in crop models beyond a simple sum of degree days is warranted. More generally, we suggest that systematically calculating a skill measure like skillT would give valuable information about the usefulness of more complex models.

### 4.4 Model averaging

As found in many studies, e-median and e-mean had prediction errors comparable to the best modeling groups. This confirmed previous evidence and theoretical considerations showing that the use of e-mean or e-median is often a good strategy (Bassu et al., 2014; Palosuo et al., 2011; Rötter et al., 2012; Wallach et al., 2018). The e-mean model is based on a simple average over simulated values, so the results from every modeling group are weighted equally. An open question in using model ensembles is whether it would be better to give more weight to models that have smaller prediction errors for the calibration data (Christensen et al., 2010), for example using Bayesian Model Averaging (Wöhling et al., 2015). The results here show that phenology predictive performance for the calibration environments is significantly correlated with predictive performance for new environments. This was also found to be the case for a study evaluating phenology prediction by modeling groups based on phenology in French environments (Wallach et al., 2019) and suggests that in these cases, it may be worthwhile to use performance-weighted model ensembles. This may be due to the fact that in these studies, the calibration and evaluation environments were similar to one another. In cases where one is extrapolating to conditions quite different than those represented by the calibration environments, performance weighting may be less useful. This once again emphasizes that it is important to define for each evaluation study whether it is an evaluation of type “interpolation” or “extrapolation”.

### 4.5 Sources of variability

A major outcome of model ensemble studies is the variability in simulated values between modeling groups, which is an indication of the uncertainty of model-based predictions (Asseng et al., 2013). Beyond a measure of the variability, it is of interest to understand the origins of the variability. One important aspect here is how differences in the model equations between model structures affect the simulated values. This however is difficult to untangle, given the multiple differences between structures. It seems that specific studies, for example modifying one specific aspect of multiple models, are needed to understand the various sources of structure uncertainty (Maiorano et al., 2016). The present study does not allow us to relate specific differences in model structure to differences in simulated results. However, it does allow us to separate two contributions to variability, namely the overall variability between model structures and the variability between different parameter values for the same model structure. An important question is the relative importance of the two, to determine priorities for reducing overall uncertainty. Parameter uncertainty can arise from uncertainty in the default values of those parameters that are fixed, from uncertainty in the choice of calibration approach (for example, the form of the objective function or the choice of parameters to estimate) and from the values of the estimated parameters, which are uncertain because there is always a limited amount of data. The within-structure variability here is a measure of the uncertainty due to choice of default values and calibration approach, but not of uncertainty in the values of the calibrated parameters. The within-structure standard deviation here is 4.3 days, compared to a between-structure standard deviation (contribution of structure) of 11.9 days. The study based on French environments found a within-structure standard deviation of 5.6 days and a between-structure standard deviation of 8.0 days (Wallach et al., 2019). Confalonieri et al. (2016) also found that the within-structure effect was in general, but not in all cases, smaller than the between-structure effect on variability.

Other studies have on the contrary focused on structural uncertainty versus uncertainty in the calibrated parameters, without taking into account uncertainty in all the default parameter values, nor uncertainty in the calibration approach chosen. Zhang et al. (2017) found that model structure explained about 80 % of the variability in simulated time to heading in rice and about 92 % of the variability in simulated time to maturity in rice, the remainder of the variability being due to parameter uncertainty. Wallach et al. (2017) found that model structure uncertainty contributed about twice as much variance as parameter uncertainty to overall simulation variance. It would be of interest to have a fuller treatment of parameter uncertainty, including both different groups using the same model structure and an estimate of the uncertainty in the parameters estimated by each group.

## 5. Conclusions

We evaluated how well 28 crop modeling groups simulate wheat phenology in Australia, in the case where both the calibration data and the evaluation data were sampled from fields in the major wheat growing areas in Australia under current climate and local management. It is important to distinguish between interpolation type prediction, as here, and extrapolation type, since they are not evaluating the same properties of modeling groups. It is also important to emphasize that evaluation concerns both model structure and parameter values, and therefore the modeling group and not just the underlying model structure. MAE for the evaluation data here ranged from 6 to 20 days depending on the modeling group, with a median of 9 days. About two thirds of the modeling groups performed better than a simple but relevant benchmark, which predicts phenology assuming a constant temperature sum for each development stage. The added complexity of crop models beyond just the effect of temperature is therefore justified in most cases. As found in many other studies, the multi-modeling group mean and median had prediction errors nearly as small as the best modeling group, suggesting that using these ensemble predictors is a good strategy. Prediction errors for calibration and evaluation environments were found to be significantly correlated, which suggests that for interpolation type studies, it would be of interest to test ensemble predictors that weight individual models based on performance for the calibration data. The variability due to modeling group for a given model structure, which reflects part of parameter uncertainty, was found to be smaller than the variability due to model structure, but was not negligible. This implies that model improvement could be achieved not only by improving model structure but also by improving parameter values.

## Acknowledgements

This work was in part supported by the Collaborative Research Center 1253 CAMPOS (Project 7: Stochastic Modelling Framework), funded by the German Research Foundation (DFG, Grant Agreement SFB 1253/1 2017), the Academy of Finland through projects AI-CropPro (316172 and 315896) and DivCSA (316215) and Natural Resources Institute Finland (Luke) through a strategic project BoostIA, the BonaRes projects ‘‘Soil3’’ (BOMA 03037514) and “I4S” (031B0513I) of the Federal Ministry of Education and Research (BMBF), Germany, the Deutsche Forschungsgemeinschaft (DFG, German Research Foundation) under Germany’s Excellence Strategy - EXC 2070 – 390732324, the project BiomassWeb of the GlobeE programme (Grant number: FKZ031A258B) funded by the Federal Ministry of Education and Research (BMBF, Germany), the EU funded SustEs project (CZ.02.1.01/0.0/0.0/16_019/0000797), the INRA ACCAF meta-programme, the German Federal Ministry of Education and Research (BMBF) in the framework of the funding measure “Soil as a Sustainable Resource for the Bioeconomy – BonaRes”, project “BonaRes (Module B): BonaRes Centre for Soil Research, subproject B” (grant 031B0511B), the National Key Research and Development Program of China (2017YFD0300205), the National Science Foundation for Distinguished Young Scholars (31725020), the Priority Academic Program Development of Jiangsu Higher Education Institutions (PAPD), the 111 project (B16026), and China Scholarship Council, the Agriculture and Agri-Food Canada’s Project 1387 under the Canadian Agricultural Partnership, the DFG Research Unit FOR 1695 ‘Agricultural Landscapes under Global Climate Change – Processes and Feedbacks on a Regional Scale, the U.S. Department of Agriculture National Institute of Food and Agriculture (award no. 2015-68007-23133) and USDA/NIFA HATCH grant N. MCL02368, the National Key Research and Development Program of China (2016YFD0300105), The Broadacre Agriculture Initiative, a research partnership between University of Southern Queensland and the Queensland Department of Agriculture and Fisheries, the JPI FACCE MACSUR2 project, funded by the Italian Ministry for Agricultural, Food, and Forestry Policies (D.M. 24064/7303/15 of 26/Nov/2015). The field work was jointly funded by CSIRO and the Grains Research and Development Corporation (GRDC) under the “Adding Value to GRDC’s National Variety Trial Network” project (CSA00027). The order in which the donors are listed is arbitrary

## SUPPLEMENTARY

**Table S1.**
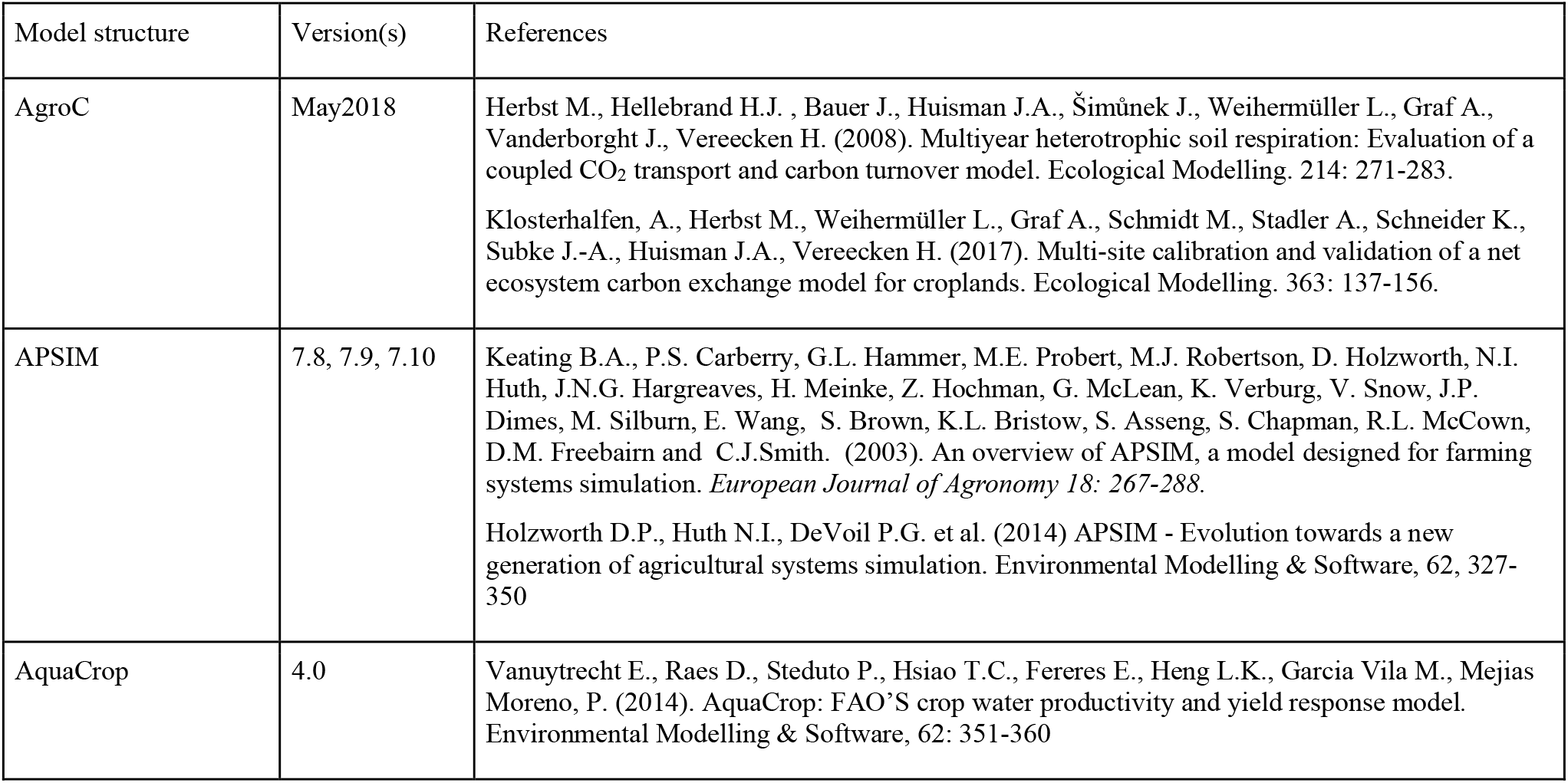
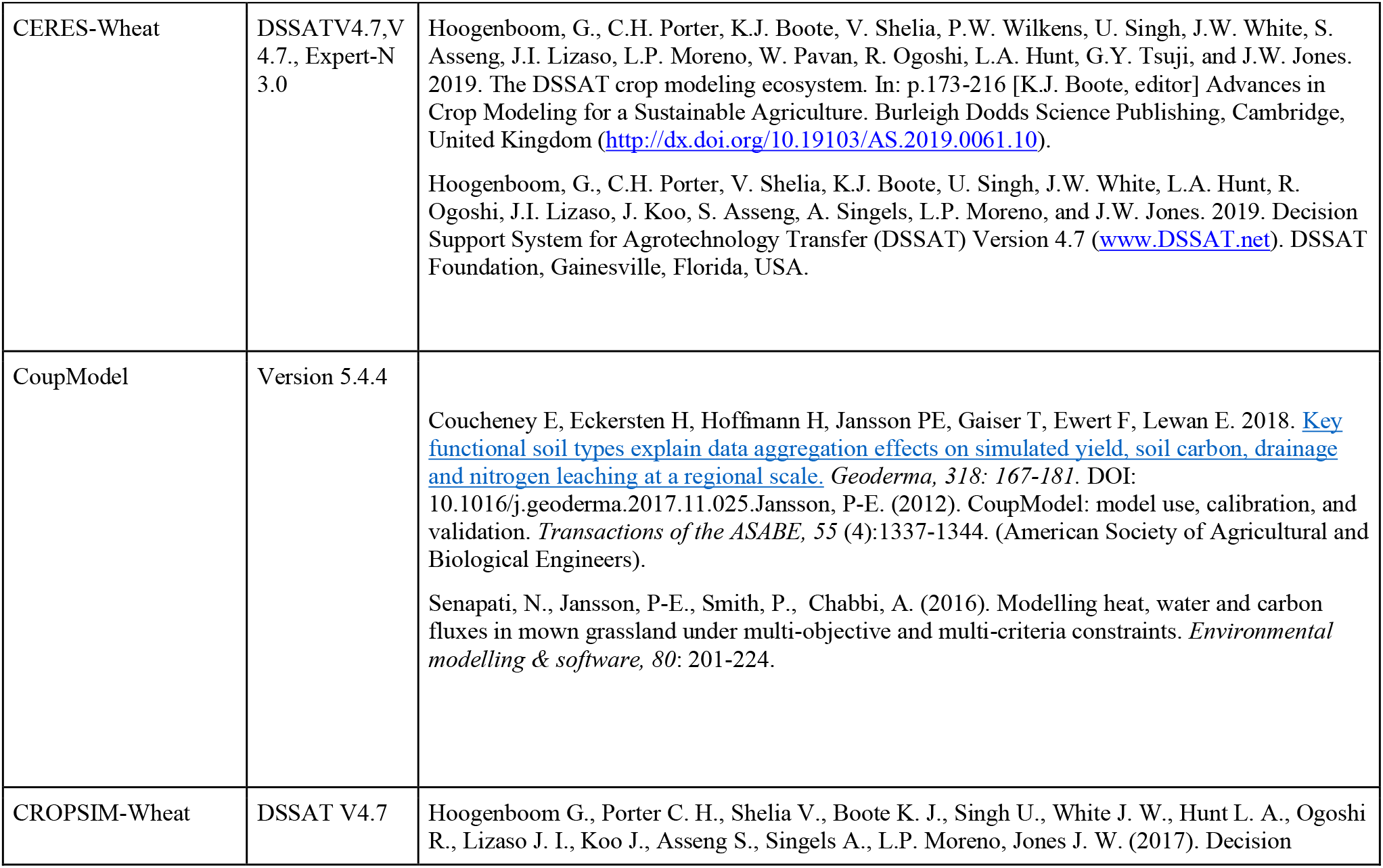
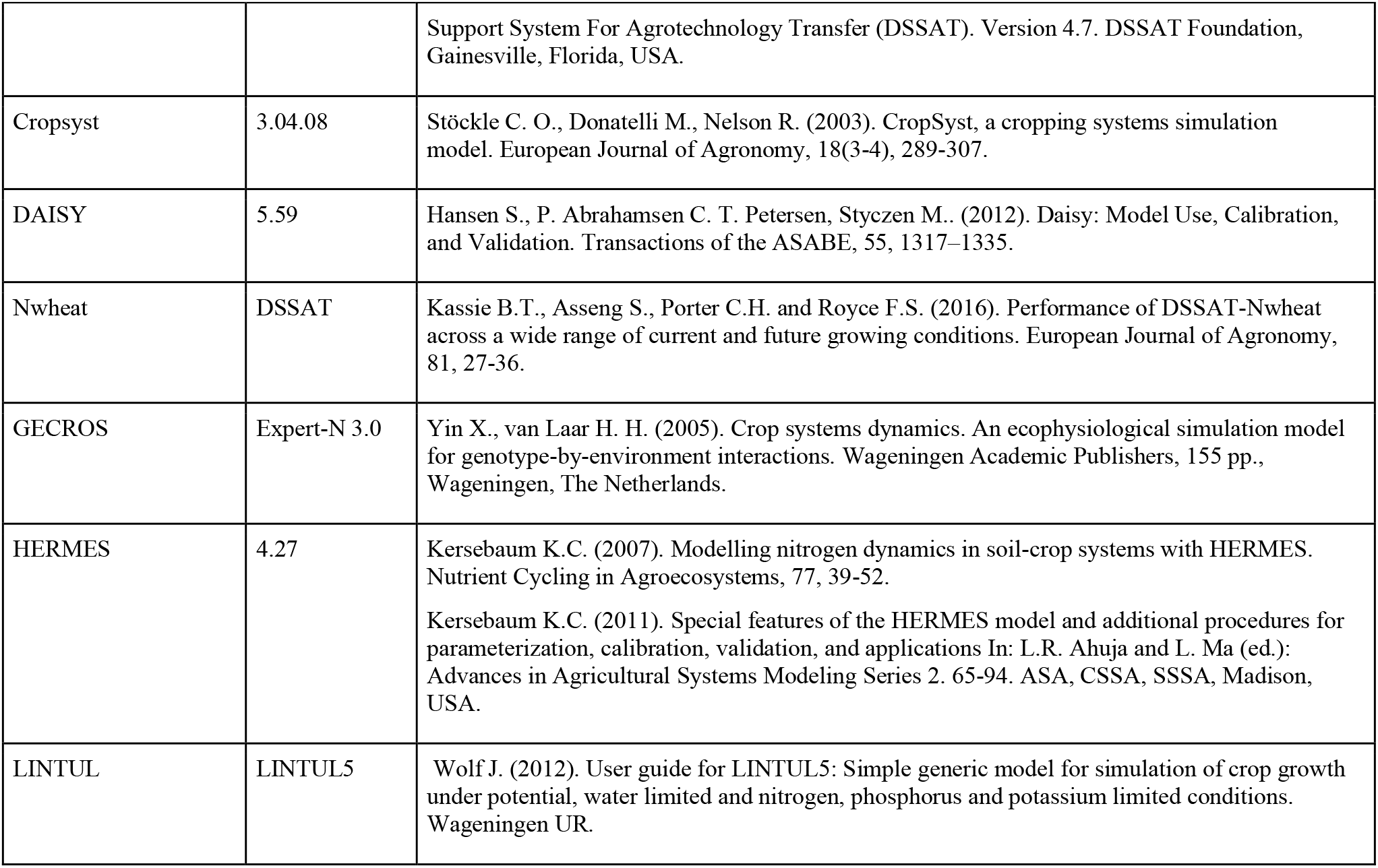
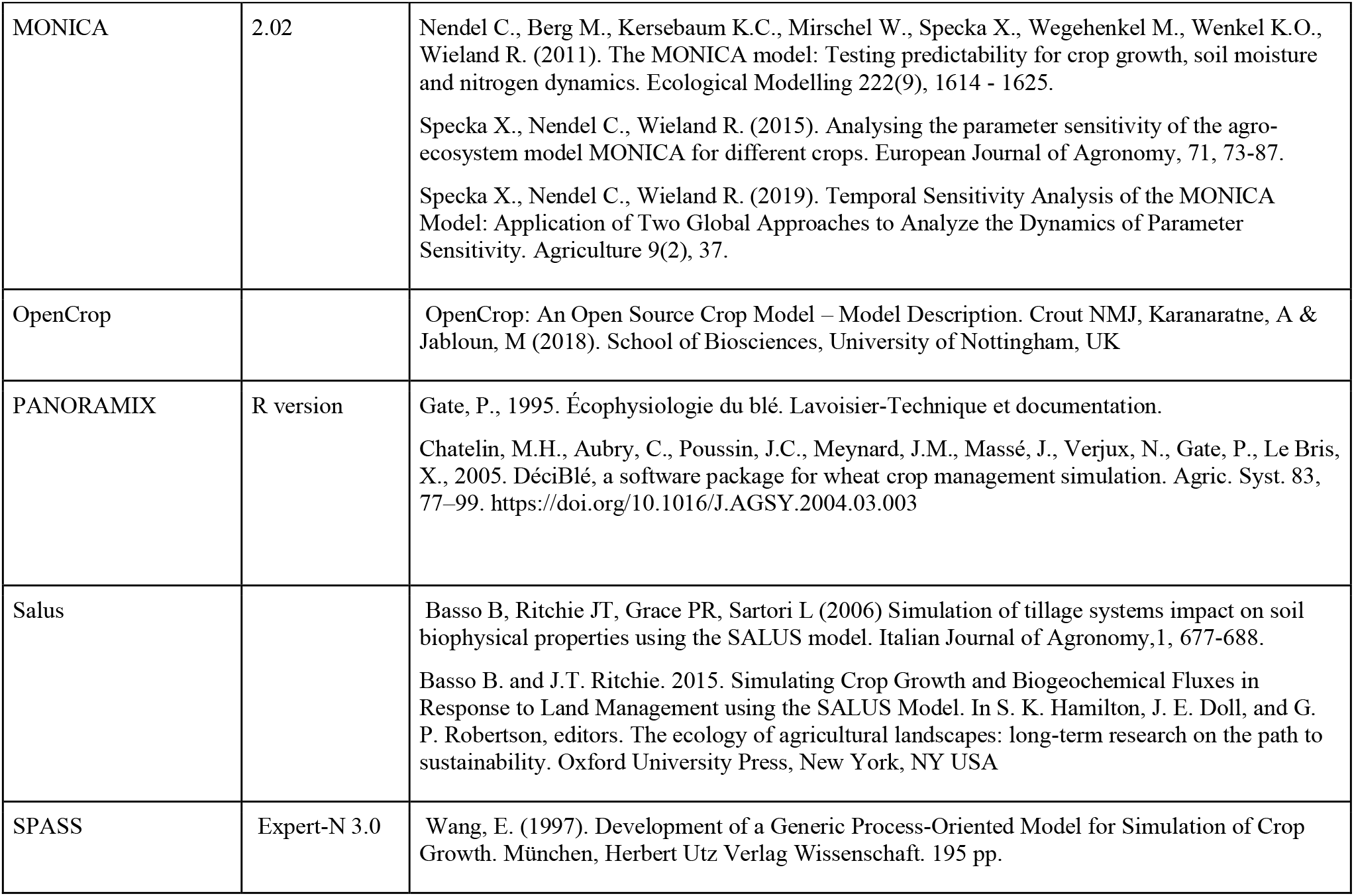
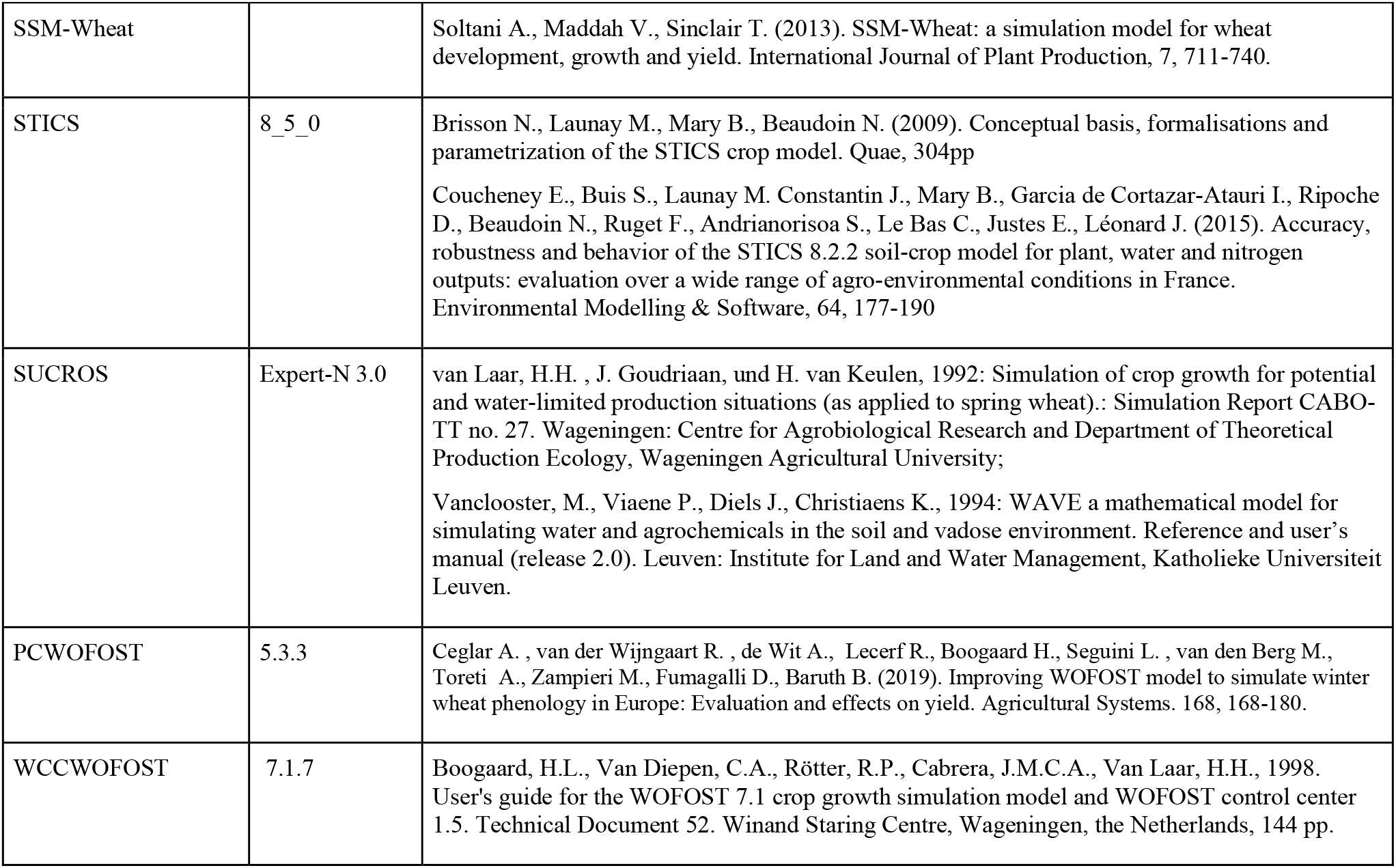
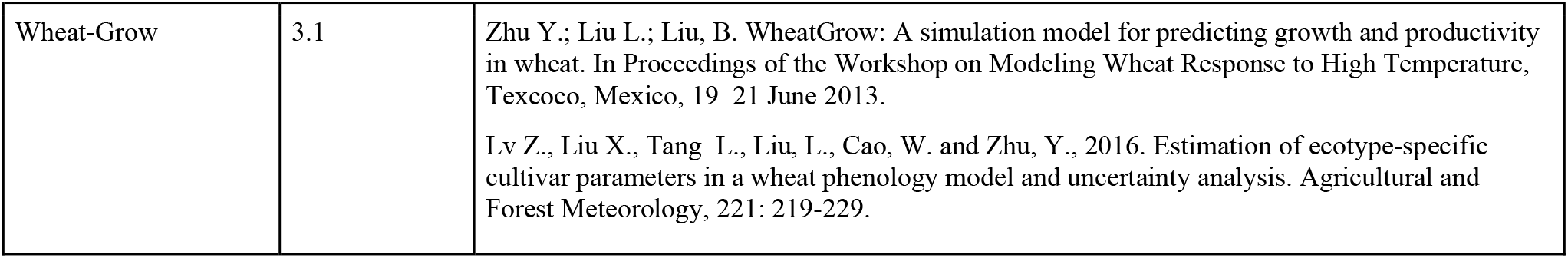
Model structures used in this study

**Figure S1.**
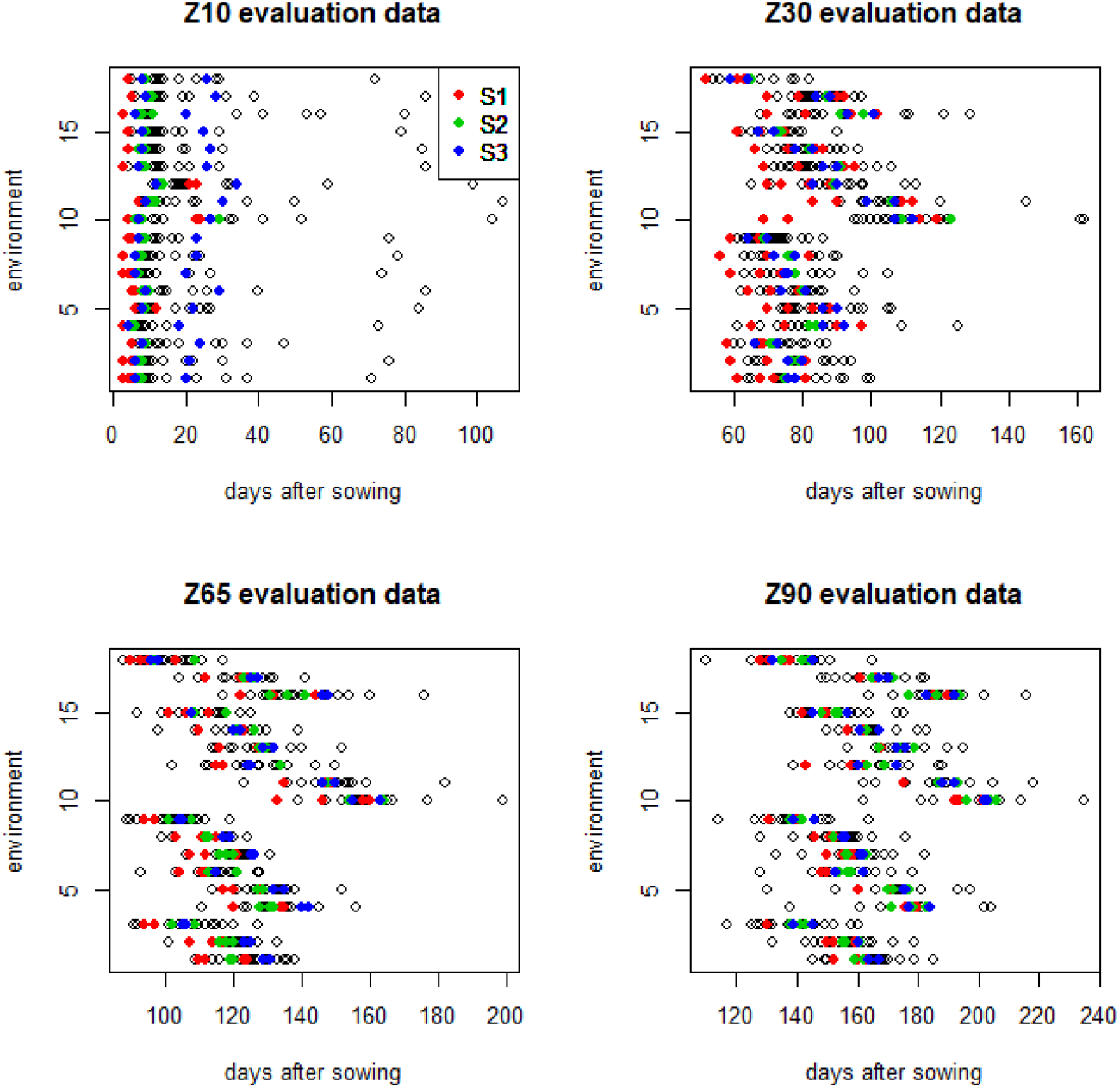
Predictions of days from sowing to Zadoks stages Z10 (emergence), Z30, Z65 and Z90 by each modeling group for each evaluation environment. Modeling groups that used the same model structure are identified by color (red for structure S1, green for structure S2, blue for structure S3).

**Table S2.**
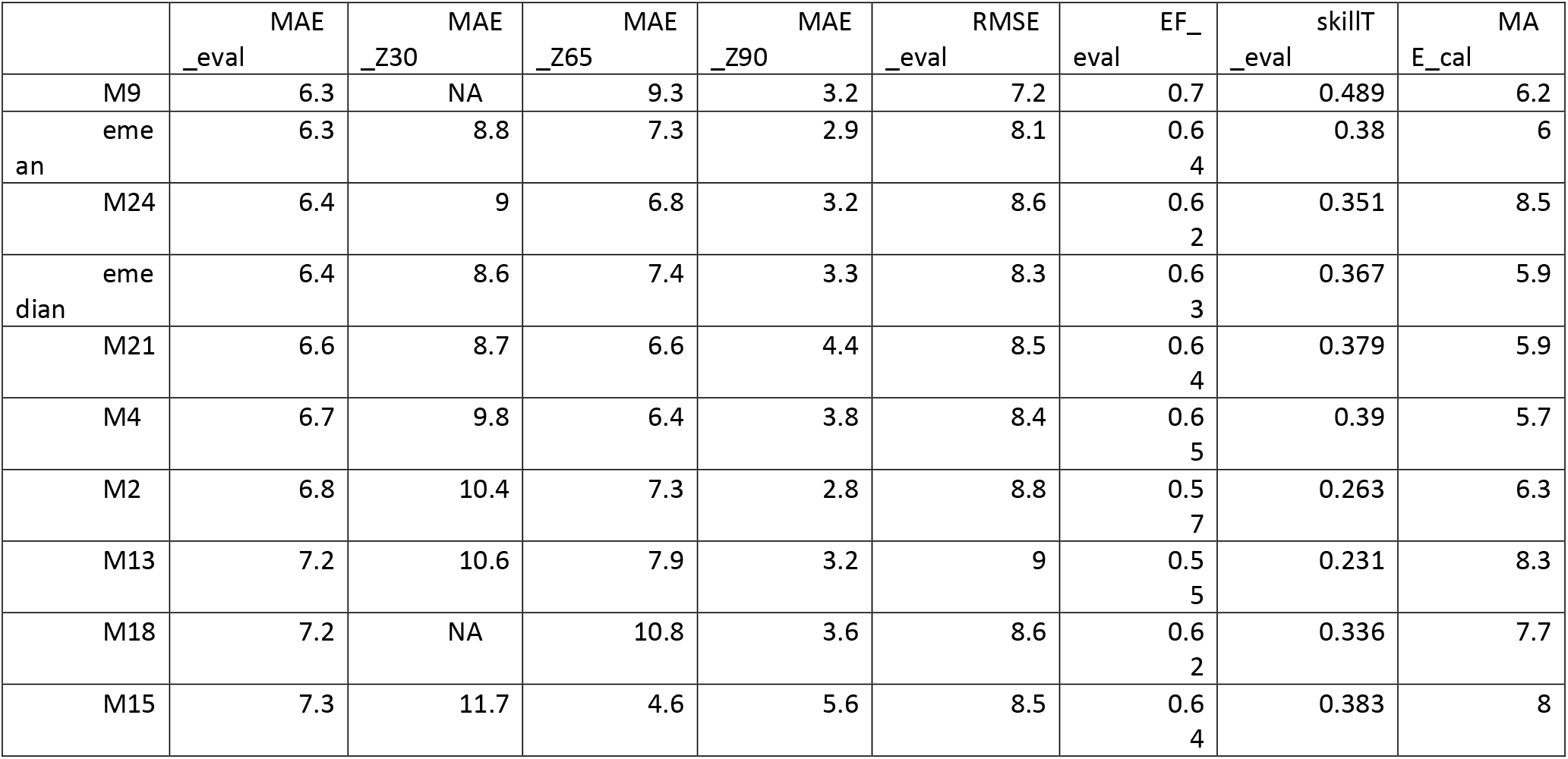
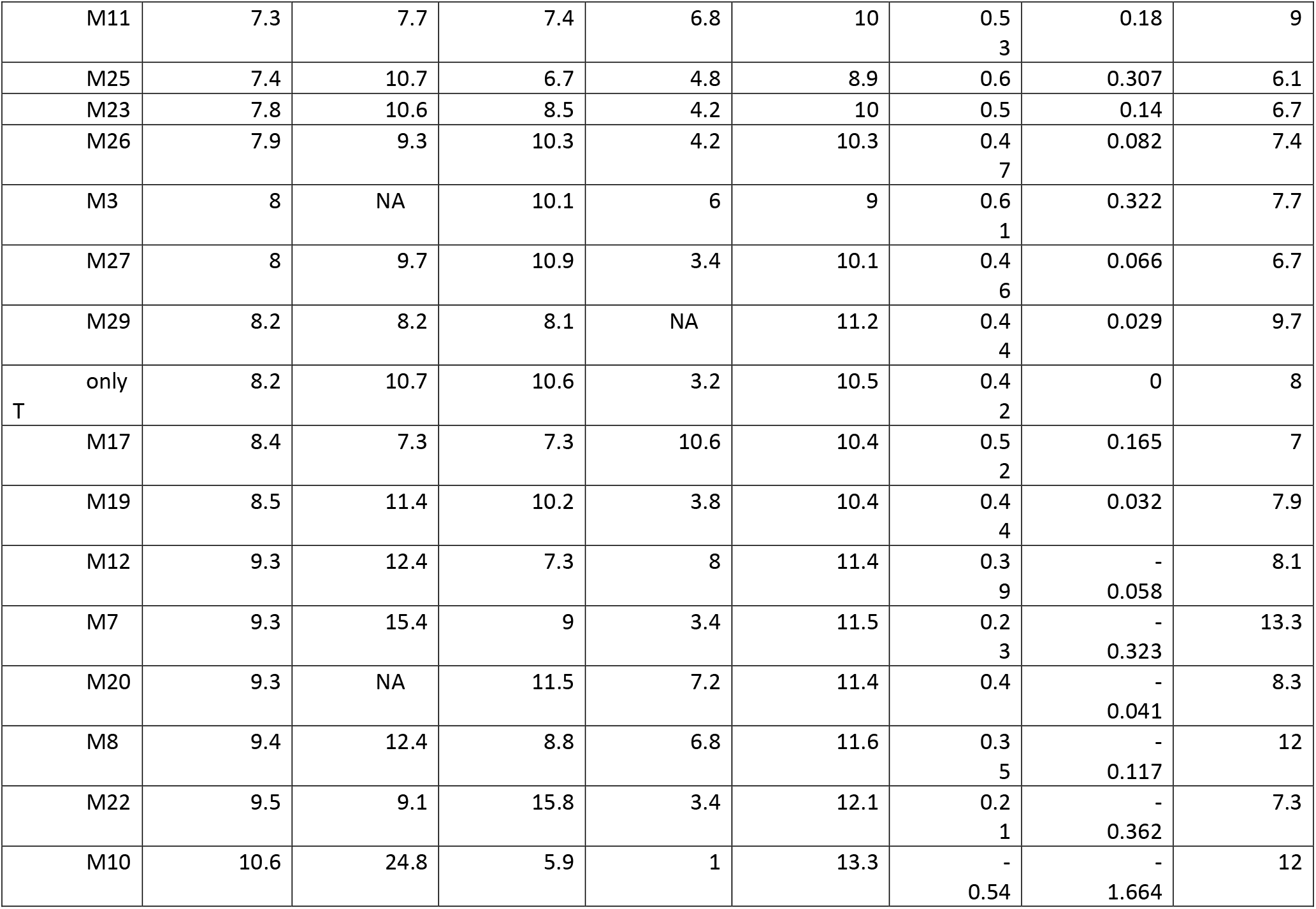
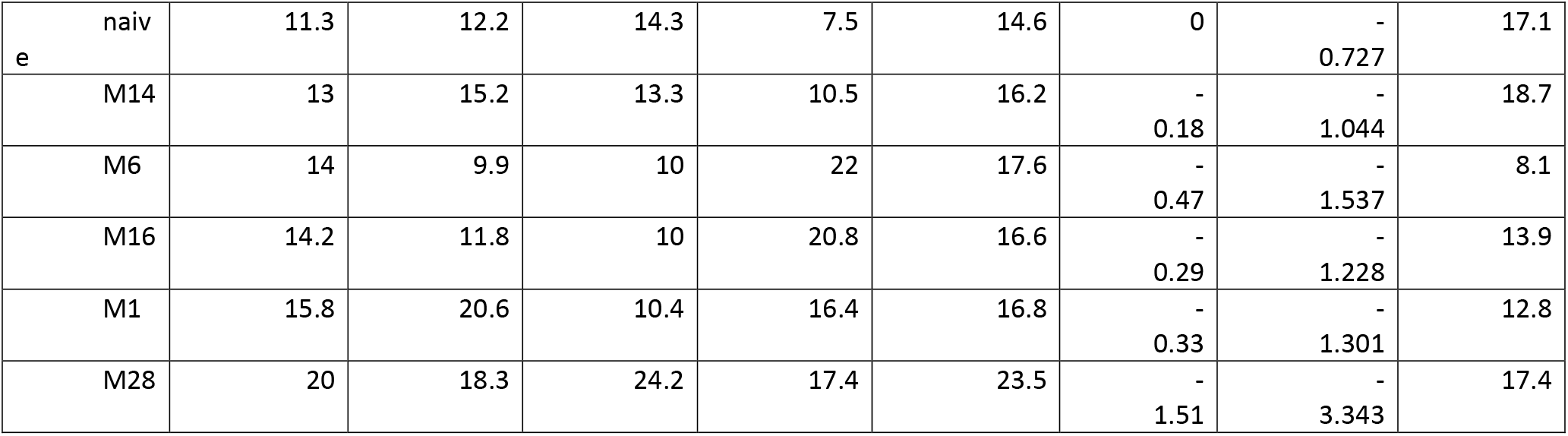
Prediction errors for each modeling group and for the models e-mean, e-median, naive and onlyT. The columns are MAE averaged over the stages Z30, Z65 and Z90 for the evaluation environments (days), MAE for each of the stages Z30, Z65 and Z90 for the evaluation environments (days), root mean squared error (RMSE) averaged over the stages Z30, Z65 and Z90 for the evaluation environments (days), the skill measures EF and skillT averaged over the stages Z30, Z65 and Z90 for the evaluation environments (unitless) and MAE averaged over the stages Z30, Z65 and Z90 for the calibration environments (days). The models are ordered by average MAE (value in first column). NA indicates that that modeling group didn’t predict the time to the indicated stage.

**Fig. S2.**
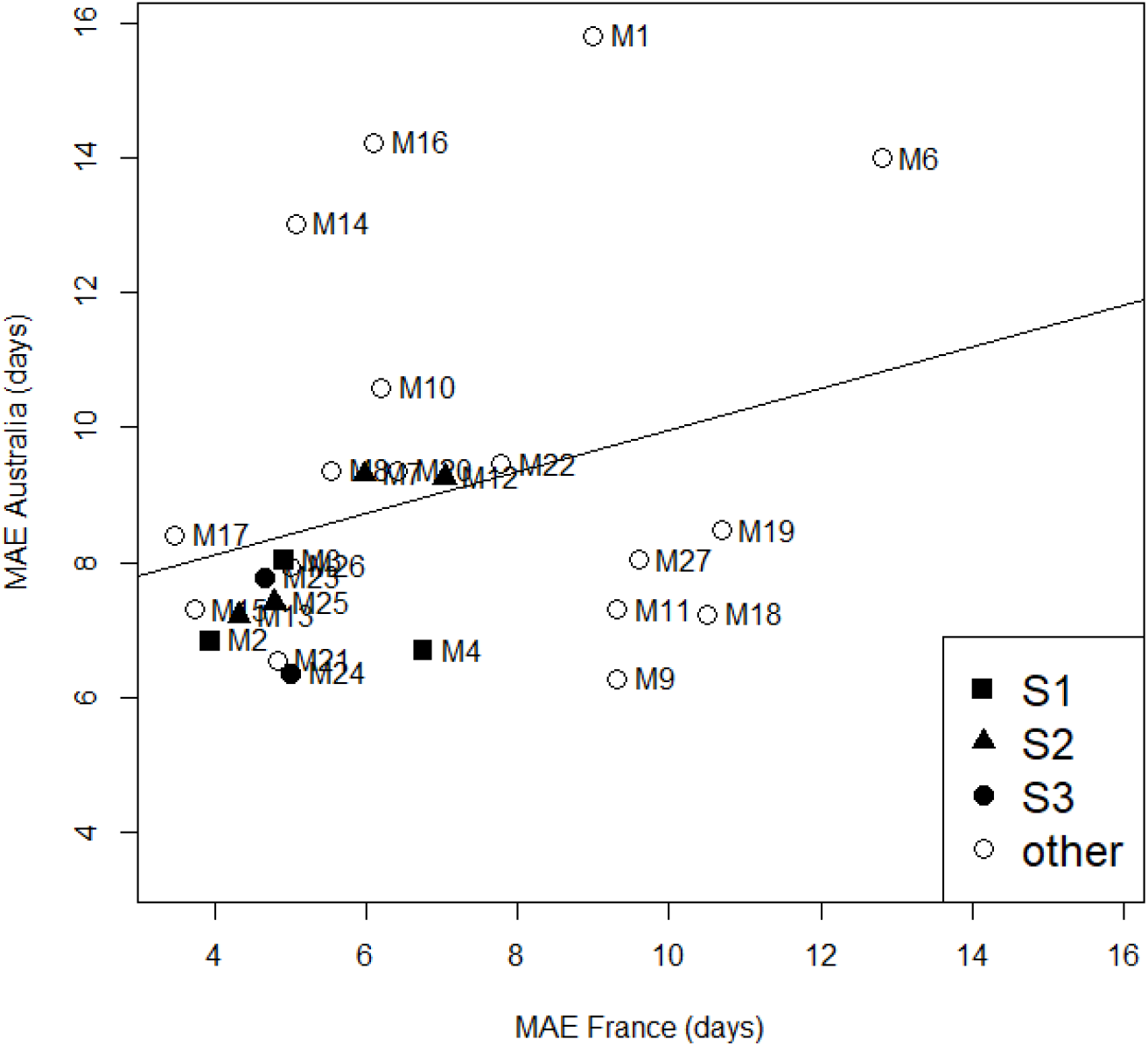
Relation between mean absolute error (MAE) for the Australian environments and MAE for the French environments, for modeling groups that participated in both studies. Values are averages over predicted development stages. Points are identified by modeling group. Modeling groups that shared the same structure (S1, S2 or S3) are identified by filled squares, triangles or circles, respectively. The regression line is y=8.23+0.24x.

## References

Andarzian, Bahram, Hoogenboom, G., Bannayan, M., Shirali, M., Andarzian, Behnam, 2015. Determining optimum sowing date of wheat using CSM-CERES-Wheat model. J. Saudi Soc. Agric. Sci. 14, 189–199. https://doi.org/10.1016/J.JSSAS.2014.04.004

Archontoulis, S. V., Miguez, F.E., Moore, K.J., 2014. A methodology and an optimization tool to calibrate phenology of short-day species included in the APSIM PLANT model: Application to soybean. Environ. Model. Softw. 62, 465–477. https://doi.org/10.1016/j.envsoft.2014.04.009

Asseng, S., Ewert, F., Rosenzweig, C., Jones, J.W., Hatfield, J.L., Ruane, A.C., Boote, K.J., Thorburn, P.J., Rötter, R.P., Cammarano, D., Brisson, N., Basso, B., Martre, P., Aggarwal, P.K., Angulo, C., Bertuzzi, P., Biernath, C., Challinor, A.J., Doltra, J., Gayler, S., Goldberg, R., Grant, R., Heng, L., Hooker, J., Hunt, L.A., Ingwersen, J., Izaurralde, R.C., Kersebaum, K.C., Müller, C., Naresh Kumar, S., Nendel, C., O’Leary, G., Olesen, J.E., Osborne, T.M., Palosuo, T., Priesack, E., Ripoche, D., Semenov, M.A., Shcherbak, I., Steduto, P., Stöckle, C., Stratonovitch, P., Streck, T., Supit, I., Tao, F., Travasso, M., Waha, K., Wallach, D., White, J.W., Williams, J.R., Wolf, J., 2013. Uncertainty in simulating wheat yields under climate change. Nat. Clim. Chang. 3, 827–832. https://doi.org/10.1038/nclimate1916

Asseng, S., Keating, B.A., Fillery, I.R.P., Gregory, P.J., Bowden, J.W., Turner, N.C., Palta, J.A., Abrecht, D.G., 2008. Performance of the APSIM-wheat model in Western Australia. F. Crop. Res. 57, 163–179.

Bao, Y., Hoogenboom, G., McClendon, R., Vellidis, G., 2017. A comparison of the performance of the CSM-CERES-Maize and EPIC models using maize variety trial data. Agric. Syst. 150, 109–119. https://doi.org/10.1016/J.AGSY.2016.10.006

Bassu, S., Brisson, N., Durand, J.-L., Boote, K., Lizaso, J., Jones, J.W., Rosenzweig, C., Ruane, A.C., Adam, M., Baron, C., Basso, B., Biernath, C., Boogaard, H., Conijn, S., Corbeels, M., Deryng, D., De Sanctis, G., Gayler, S., Grassini, P., Hatfield, J., Hoek, S., Izaurralde, C., Jongschaap, R., Kemanian, A.R., Kersebaum, K.C., Kim, S.-H., Kumar, N.S., Makowski, D., Müller, C., Nendel, C., Priesack, E., Pravia, M.V., Sau, F., Shcherbak, I., Tao, F., Teixeira, E., Timlin, D., Waha, K., 2014. How do various maize crop models vary in their responses to climate change factors? Glob. Chang. Biol. 20, 2301–20. https://doi.org/10.1111/gcb.12520

Biernath, C., Gayler, S., Bittner, S., Klein, C., Högy, P., Fangmeier, A., Priesack, E., 2011. Evaluating the ability of four crop models to predict different environmental impacts on spring wheat grown in open-top chambers. Eur. J. Agron. 35, 71–82. https://doi.org/10.1016/j.eja.2011.04.001

Boote, K.J., Jones, J.W., Hoogenboom, G., 2008. Crop simulation models as tools for agro-advisories for weather and disease effects on production. J. Agrometeorol. 10, 9–17.

Boote, K.J., Jones, J.W., Hoogenboom, G., White, J.W., 2010. The Role of Crop Systems Simulation in Agriculture and Environment. Int. J. Agric. Environ. Inf. Syst. 1, 41–54.

Casella, G., Berger, R.L., 1990. Statistical Inference. Wadsworth and Brooks, Pacific Grove, CA.

Ceglar, A., van der Wijngaart, R., de Wit, A., Lecerf, R., Boogaard, H., Seguini, L., van den Berg, M., Toreti, A., Zampieri, M., Fumagalli, D., Baruth, B., 2019. Improving WOFOST model to simulate winter wheat phenology in Europe: Evaluation and effects on yield. Agric. Syst. 168, 168–180. https://doi.org/10.1016/J.AGSY.2018.05.002

Chauhan, Y.S., Ryan, M., Chandra, S., Sadras, V.O., 2019. Accounting for soil moisture improves prediction of flowering time in chickpea and wheat. Sci. Rep. 9, 7510. https://doi.org/10.1038/s41598-019-43848-6

Christensen, J., Kjellström, E., Giorgi, F., Lenderink, G., Rummukainen, M., 2010. Weight assignment in regional climate models. Clim. Res. 44, 179–194. https://doi.org/10.3354/cr00916

Confalonieri, R., Orlando, F., Paleari, L., Stella, T., Gilardelli, C., Movedi, E., Pagani, V., Cappelli, G., Vertemara, A., Alberti, L., Alberti, P., Atanassiu, S., Bonaiti, M., Cappelletti, G., Ceruti, M., Confalonieri, A., Corgatelli, G., Corti, P., Dell’Oro, M., Ghidoni, A., Lamarta, A., Maghini, A., Mambretti, M., Manchia, A., Massoni, G., Mutti, P., Pariani, S., Pasini, D., Pesenti, A., Pizzamiglio, G., Ravasio, A., Rea, A., Santorsola, D., Serafini, G., Slavazza, M., Acutis, M., 2016. Uncertainty in crop model predictions: What is the role of users? Environ. Model. Softw. 81, 165–173. https://doi.org/10.1016/j.envsoft.2016.04.009

Corripio, J.G., n.d. insol: Solar Radiation. R package version 1.2. 2019.

Efron, B., 1986. How Biased is the Apparent Error Rate of a Prediction Rule? J. Am. Stat. Assoc. 81, 461–470. https://doi.org/10.1080/01621459.1986.10478291

Flohr, B.M., Hunt, J.R., Kirkegaard, J.A., Evans, J.R., 2017. Water and temperature stress define the optimal flowering period for wheat in south-eastern Australia. F. Crop. Res. v. 209, 108–119. https://doi.org/10.1016/j.fcr.2017.04.012

Hussain, J., Khaliq, T., Ahmad, A., Akhtar, J., 2018. Performance of four crop model for simulations of wheat phenology, leaf growth, biomass and yield across planting dates. PLoS One 13, e0197546. https://doi.org/10.1371/journal.pone.0197546

Johnen, T., Boettcher, U., Kage, H., 2012. A variable thermal time of the double ridge to flag leaf emergence phase improves the predictive quality of a CERES-Wheat type phenology model. Comput. Electron. Agric. 89, 62–69. https://doi.org/10.1016/J.COMPAG.2012.08.002

Kumudini, S., Andrade, F.H., Boote, K.J., Brown, G.A., Dzotsi, K.A., Edmeades, G.O., Gocken, T., Goodwin, M., Halter, A.L., Hammer, G.L., Hatfield, J.L., Jones, J.W., Kemanian, A.R., Kim, S.-H., Kiniry, J., Lizaso, J.I., Nendel, C., Nielsen, R.L., Parent, B., Stöckle, C.O., Tardieu, F., Thomison, P.R., Timlin, D.J., Vyn, T.J., Wallach, D., Yang, H.S., Tollenaar, M., 2014. Predicting maize phenology: Intercomparison of functions for developmental response to temperature. Agron. J. 106, 2087–2097. https://doi.org/10.2134/agronj14.0200

Lawes, R.A., Huth, N.D., Hochman, Z., 2016. Commercially available wheat cultivars are broadly adapted to location and time of sowing in Australia’s grain zone. Eur. J. Agron. 77, 38–46. https://doi.org/10.1016/J.EJA.2016.03.009

Lindstrom, M.J., Papendick, R.I., Koehler, F.E., 1976. A Model to Predict Winter Wheat Emergence as Affected by Soil Temperature, Water Potential, and Depth of Planting1. Agron. J. 68, 137–141. https://doi.org/10.2134/agronj1976.00021962006800010038x

Luo, Q., O’Leary, G., Cleverly, J., Eamus, D., 2018. Effectiveness of time of sowing and cultivar choice for managing climate change: wheat crop phenology and water use efficiency. Int. J. Biometeorol. 62, 1049–1061. https://doi.org/10.1007/s00484-018-1508-4

Maiorano, A., Martre, P., Asseng, S., Ewert, F., Müller, C., Rötter, R.P., Ruane, A.C., Semenov, M.A., Wallach, D., Wang, E., Alderman, P.D., Kassie, B.T., Biernath, C., Basso, B., Cammarano, D., Challinor, A.J., Doltra, J., Dumont, B., Rezaei, E.E., Gayler, S., Kersebaum, K.C., Kimball, B.A., Koehler, A.-K., Liu, B., O’Leary, G.J., Olesen, J.E., Ottman, M.J., Priesack, E., Reynolds, M., Stratonovitch, P., Streck, T., Thorburn, P.J., Waha, K., Wall, G.W., White, J.W., Zhao, Z., Zhu, Y., 2016. Crop model improvement reduces the uncertainty of the response to temperature of multi-model ensembles. F. Crop. Res. https://doi.org/10.1016/j.fcr.2016.05.001

McCuen, R.H., Knight, Z., Cutter, A.G., 2006. Evaluation of the Nash–Sutcliffe Efficiency Index. J. Hydrol. Eng. 11, 597–602. https://doi.org/10.1061/(ASCE)1084-0699(2006)11:6(597)

Palosuo, T., Kersebaum, K.C., Angulo, C., Hlavinka, P., Moriondo, M., Olesen, J.E., Patil, R.H., Ruget, F., Rumbaur, C., Takáč, J., Trnka, M., Bindi, M., Çaldağ, B., Ewert, F., Ferrise, R., Mirschel, W., Şaylan, L., Šiška, B., Rötter, R., 2011. Simulation of winter wheat yield and its variability in different climates of Europe: A comparison of eight crop growth models. Eur. J. Agron. 35, 103–114. https://doi.org/10.1016/j.eja.2011.05.001

R Core Team, 2017. A language and Environment for Statistical Computing.

Rötter, R.P., Palosuo, T., Kersebaum, K.C., Angulo, C., Bindi, M., Ewert, F., Ferrise, R., Hlavinka, P., Moriondo, M., Nendel, C., Olesen, J.E., Patil, R.H., Ruget, F., Takáč, J., Trnka, M., 2012. Simulation of spring barley yield in different climatic zones of Northern and Central Europe: A comparison of nine crop models. F. Crop. Res. 133, 23–36. https://doi.org/10.1016/j.fcr.2012.03.016

Sadras, V.O., Monzon, J.P., 2006. Modelled wheat phenology captures rising temperature trends: Shortened time to flowering and maturity in Australia and Argentina. F. Crop. Res. 99, 136–146. https://doi.org/10.1016/J.FCR.2006.04.003

Teluguntla, P., Thenkabail, P.S., Oliphant, A., Xiong, J., Gumma, M.K., Congalton, R.G., Yadav, K., Huete, A., 2018. A 30-m landsat-derived cropland extent product of Australia and China using random forest machine learning algorithm on Google Earth Engine cloud computing platform. ISPRS J. Photogramm. Remote Sens. 144, 325–340. https://doi.org/10.1016/J.ISPRSJPRS.2018.07.017

van Bussel, L.G.J., Stehfest, E., Siebert, S., Müller, C., Ewert, F., 2015. Simulation of the phenological development of wheat and maize at the global scale. Glob. Ecol. Biogeogr. 24, 1018–1029. https://doi.org/10.1111/geb.12351

Wallach, D., Martre, P., Liu, B., Asseng, S., Ewert, F., Thorburn, P.J., van Ittersum, M., Aggarwal, P.K., Ahmed, M., Basso, B., Biernath, C., Cammarano, D., Challinor, A.J., De Sanctis, G., Dumont, B., Eyshi Rezaei, E., Fereres, E., Fitzgerald, G.J., Gao, Y., Garcia-Vila, M., Gayler, S., Girousse, C., Hoogenboom, G., Horan, H., Izaurralde, R.C., Jones, C.D., Kassie, B.T., Kersebaum, K.C., Klein, C., Koehler, A.-K., Maiorano, A., Minoli, S., Müller, C., Naresh Kumar, S., Nendel, C., O’Leary, G.J., Palosuo, T., Priesack, E., Ripoche, D., Rötter, R.P., Semenov, M.A., Stöckle, C., Stratonovitch, P., Streck, T., Supit, I., Tao, F., Wolf, J., Zhang, Z., 2018. Multimodel ensembles improve predictions of crop-environment-management interactions. Glob. Chang. Biol. 24, 5072–5083. https://doi.org/10.1111/gcb.14411

Wallach, D., Nissanka, S.P., Karunaratne, A.S., Weerakoon, W.M.W., Thorburn, P.J., Boote, K.J., Jones, J.W., 2017. Accounting for both parameter and model structure uncertainty in crop model predictions of phenology: A case study on rice. Eur. J. Agron. 88. https://doi.org/10.1016/j.eja.2016.05.013

Wallach, D., Palosuo, T., Thorburn, P., Seidel, S.J., Gourdain, E., Asseng, S., Basso, B., Buis, S., Crout, N.M.J., Dibari, C., Dumont, B., Ferrise, R., Gaiser, T., Garcia, C., Gayler, S., Ghahramani, A., Hochman, Z., Hoek, S., Horan, H., Hoogenboom, G., Huang, M., Jabloun, M., Jing, Q., Justes, E., Kersebaum, K.C., Klosterhalfen, A., Launay, M., Luo, Q., Maestrini, B., Mielenz, H., Moriondo, M., Nariman Zadeh, H., Olesen, J.E., Poyda, A., Priesack, E., Pullens, J.W.M., Qian, B., Schütze, N., Shelia, V., Souissi, A., Specka, X., Srivastava, A.K., Stella, T., Streck, T., Trombi, G., Wallor, E., Wang, J., Weber, T.K.D., Weihermüller, L., de Wit, A., Wöhling, T., Xiao, L., Zhao, C., Zhu, Y., 2019. How well do crop models predict phenology, given calibration data from the target population? bioRxiv 708578. https://doi.org/10.1101/708578

Wang, E., Martre, P., Zhao, Z., Ewert, F., Maiorano, A., Rötter, R.P., Kimball, B.A., Ottman, M.J., Wall, G.W., White, J.W., Reynolds, M.P., Alderman, P.D., Aggarwal, P.K., Anothai, J., Basso, B., Biernath, C., Cammarano, D., Challinor, A.J., De Sanctis, G., Doltra, J., Fereres, E., Garcia-Vila, M., Gayler, S., Hoogenboom, G., Hunt, L.A., Izaurralde, R.C., Jabloun, M., Jones, C.D., Kersebaum, K.C., Koehler, A.-K., Liu, L., Müller, C., Naresh Kumar, S., Nendel, C., O’Leary, G., Olesen, J.E., Palosuo, T., Priesack, E., Eyshi Rezaei, E., Ripoche, D., Ruane, A.C., Semenov, M.A., Shcherbak, I., Stöckle, C., Stratonovitch, P., Streck, T., Supit, I., Tao, F., Thorburn, P., Waha, K., Wallach, D., Wang, Z., Wolf, J., Zhu, Y., Asseng, S., 2017. The uncertainty of crop yield projections is reduced by improved temperature response functions. Nat. Plants 3, 1–13. https://doi.org/10.1038/nplants.2017.102

Wang, H., Cutforth, H., McCaig, T., McLeod, G., Brandt, K., Lemke, R., Goddard, T., Sprout, C., 2009. Predicting the time to 50% seedling emergence in wheat using a Beta model. NJAS - Wageningen J. Life Sci. 57, 65–71. https://doi.org/https://doi.org/10.1016/j.njas.2009.07.003

Willmott, C.J., Matsuura, K., 2005. Advantages of the mean absolute error (MAE) over the root mean square error (RMSE) in assessing average model performance. Clim. Res. 30, 79–82.

Wöhling, T., Schöniger, A., Gayler, S., Nowak, W., 2015. Bayesian model averaging to explore the worth of data for soil-plant model selection and prediction. Water Resour. Res. 51, 2825–2846. https://doi.org/10.1002/2014WR016292

Workman, D., 2020. Worldstopexports [WWW Document]. URL http://www.worldstopexports.com/wheat-exports-country/ (accessed 3.10.20).

Yuan, S., Peng, S., Li, T., 2017. Evaluation and application of the ORYZA rice model under different crop managements with high-yielding rice cultivars in central China. F. Crop. Res. 212, 115–125. https://doi.org/10.1016/J.FCR.2017.07.010

Zadoks, J.C., Chzang, T.T., Konzak, C.F., 1974. A decimal code for the growth stages of cereals. Weed Res. 14, 415–421. https://doi.org/10.1111/j.1365-3180.1974.tb01084.x

Zhang, S., Tao, F., Zhang, Z., 2017. Uncertainty from model structure is larger than that from model parameters in simulating rice phenology in China. Eur. J. Agron. 87, 30–39. https://doi.org/10.1016/j.eja.2017.04.004

