## Supplementary Material for "Multi-model evaluation of phenology prediction for wheat in Australia"

| Model structure | Version(s) | References |
|---|---|---|
| AgroC | May2018 | Herbst M., Hellebrand H.J. , Bauer J., Huisman J.A., Šimůnek J., Weihermüller L., Graf A., Vanderborght J., Vereecken H. (2008). Multiyear heterotrophic soil respiration: Evaluation of a coupled $CO_2$ transport and carbon turnover model. Ecological Modelling. 214: 271-283.<br><br>Klosterhalfen, A., Herbst M., Weihermüller L., Graf A., Schmidt M., Stadler A., Schneider K., Subke J.-A., Huisman J.A., Vereecken H. (2017). Multi-site calibration and validation of a net ecosystem carbon exchange model for croplands. Ecological Modelling. 363: 137-156. |
| APSIM | 7.8, 7.9, 7.10 | Keating B.A., P.S. Carberry, G.L. Hammer, M.E. Probert, M.J. Robertson, D. Holzworth, N.I. Huth, J.N.G. Hargreaves, H. Meinke, Z. Hochman, G. McLean, K. Verburg, V. Snow, J.P. Dimes, M. Silburn, E. Wang, S. Brown, K.L. Bristow, S. Asseng, S. Chapman, R.L. McCown, D.M. Freebairn and C.J.Smith. (2003). An overview of APSIM, a model designed for farming systems simulation. *European Journal of Agronomy 18: 267-288.*<br><br>Holzworth D.P., Huth N.I., DeVoil P.G. et al. (2014) APSIM - Evolution towards a new generation of agricultural systems simulation. Environmental Modelling & Software, 62, 327-350 |
| AquaCrop | 4.0 | Vanuytrecht E., Raes D., Steduto P., Hsiao T.C., Fereres E., Heng L.K., Garcia Vila M., Mejias Moreno, P. (2014). AquaCrop: FAO'S crop water productivity and yield response model. Environmental Modelling & Software, 62: 351-360 |
| CERES-Wheat | DSSATV4.7,V 4.7., Expert-N 3.0 | Hoogenboom, G., C.H. Porter, K.J. Boote, V. Shelia, P.W. Wilkens, U. Singh, J.W. White, S. Asseng, J.I. Lizaso, L.P. Moreno, W. Pavan, R. Ogoshi, L.A. Hunt, G.Y. Tsuji, and J.W. Jones. 2019. The DSSAT crop modeling ecosystem. In: p.173-216 [K.J. Boote, editor] Advances in Crop Modeling for a Sustainable Agriculture. Burleigh Dodds Science Publishing, Cambridge, United Kingdom (http://dx.doi.org/10.19103/AS.2019.0061.10). |

| | | |
|---|---|---|
| | | Hoogenboom, G., C.H. Porter, V. Shelia, K.J. Boote, U. Singh, J.W. White, L.A. Hunt, R. Ogoshi, J.I. Lizaso, J. Koo, S. Asseng, A. Singels, L.P. Moreno, and J.W. Jones. 2019. Decision Support System for Agrotechnology Transfer (DSSAT) Version 4.7 (www.DSSAT.net). DSSAT Foundation, Gainesville, Florida, USA. |
| CoupModel | Version 5.4.4 | Coucheney E, Eckersten H, Hoffmann H, Jansson PE, Gaiser T, Ewert F, Lewan E. 2018. Key functional soil types explain data aggregation effects on simulated yield, soil carbon, drainage and nitrogen leaching at a regional scale. *Geoderma*, *318: 167-181*. DOI: 10.1016/j.geoderma.2017.11.025.Jansson, P-E. (2012). CoupModel: model use, calibration, and validation. *Transactions of the ASABE*, *55* (4):1337-1344. (American Society of Agricultural and Biological Engineers).<br><br>Senapati, N., Jansson, P-E., Smith, P., Chabbi, A. (2016). Modelling heat, water and carbon fluxes in mown grassland under multi-objective and multi-criteria constraints. *Environmental modelling & software*, *80*: 201-224. |
| CROPSIM-Wheat | DSSAT V4.7 | Hoogenboom G., Porter C. H., Shelia V., Boote K. J., Singh U., White J. W., Hunt L. A., Ogoshi R., Lizaso J. I., Koo J., Asseng S., Singels A., L.P. Moreno, Jones J. W. (2017). Decision Support System For Agrotechnology Transfer (DSSAT). Version 4.7. DSSAT Foundation, Gainesville, Florida, USA. |
| Cropsyst | 3.04.08 | Stöckle C. O., Donatelli M., Nelson R. (2003). CropSyst, a cropping systems simulation model. European Journal of Agronomy, 18(3-4), 289-307. |

| | | |
|---|---|---|
| DAISY | 5.59 | Hansen S., P. Abrahamsen C. T. Petersen, Styczen M.. (2012). Daisy: Model Use, Calibration, and Validation. Transactions of the ASABE, 55, 1317–1335. |
| Nwheat | DSSAT | Kassie B.T., Asseng S., Porter C.H. and Royce F.S. (2016). Performance of DSSAT-Nwheat across a wide range of current and future growing conditions. European Journal of Agronomy, 81, 27-36. |
| GECROS | Expert-N 3.0 | Yin X., van Laar H. H. (2005). Crop systems dynamics. An ecophysiological simulation model for genotype-by-environment interactions. Wageningen Academic Publishers, 155 pp., Wageningen, The Netherlands. |
| HERMES | 4.27 | Kersebaum K.C. (2007). Modelling nitrogen dynamics in soil-crop systems with HERMES. Nutrient Cycling in Agroecosystems, 77, 39-52.<br><br>Kersebaum K.C. (2011). Special features of the HERMES model and additional procedures for parameterization, calibration, validation, and applications In: L.R. Ahuja and L. Ma (ed.): Advances in Agricultural Systems Modeling Series 2. 65-94. ASA, CSSA, SSSA, Madison, USA. |
| LINTUL | LINTUL5 | Wolf J. (2012). User guide for LINTUL5: Simple generic model for simulation of crop growth under potential, water limited and nitrogen, phosphorus and potassium limited conditions. Wageningen UR. |
| MONICA | 2.02 | Nendel C., Berg M., Kersebaum K.C., Mirschel W., Specka X., Wegehenkel M., Wenkel K.O., Wieland R. (2011). The MONICA model: Testing predictability for crop growth, soil moisture and nitrogen dynamics. Ecological Modelling 222(9), 1614 - 1625.<br><br>Specka X., Nendel C., Wieland R. (2015). Analysing the parameter sensitivity of the agro-ecosystem model MONICA for different crops. European Journal of Agronomy, 71, 73-87. |

| | | |
|---|---|---|
| | | Specka X., Nendel C., Wieland R. (2019). Temporal Sensitivity Analysis of the MONICA Model: Application of Two Global Approaches to Analyze the Dynamics of Parameter Sensitivity. Agriculture 9(2), 37. |
| OpenCrop | | OpenCrop: An Open Source Crop Model – Model Description. Crout NMJ, Karanaratne, A & Jabloun, M (2018). School of Biosciences, University of Nottingham, UK |
| PANORAMIX | R version | Gate, P., 1995. Écophysiologie du blé. Lavoisier-Technique et documentation.<br><br>Chatelin, M.H., Aubry, C., Poussin, J.C., Meynard, J.M., Massé, J., Verjux, N., Gate, P., Le Bris, X., 2005. DéciBlé, a software package for wheat crop management simulation. Agric. Syst. 83, 77–99. https://doi.org/10.1016/J.AGSY.2004.03.003 |
| Salus | | Basso B, Ritchie JT, Grace PR, Sartori L (2006) Simulation of tillage systems impact on soil biophysical properties using the SALUS model. Italian Journal of Agronomy,1, 677-688.<br><br>Basso B. and J.T. Ritchie. 2015. Simulating Crop Growth and Biogeochemical Fluxes in Response to Land Management using the SALUS Model. In S. K. Hamilton, J. E. Doll, and G. P. Robertson, editors. The ecology of agricultural landscapes: long-term research on the path to sustainability. Oxford University Press, New York, NY USA |
| SPASS | Expert-N 3.0 | Wang, E. (1997). Development of a Generic Process-Oriented Model for Simulation of Crop Growth. München, Herbert Utz Verlag Wissenschaft. 195 pp. |
| SSM-Wheat | | Soltani A., Maddah V., Sinclair T. (2013). SSM-Wheat: a simulation model for wheat development, growth and yield. International Journal of Plant Production, 7, 711-740. |

| | | |
|---|---|---|
| STICS | 8_5_0 | Brisson N., Launay M., Mary B., Beaudoin N. (2009). Conceptual basis, formalisations and parametrization of the STICS crop model. Quae, 304pp<br><br>Coucheney E., Buis S., Launay M. Constantin J., Mary B., Garcia de Cortazar-Atauri I., Ripoche D., Beaudoin N., Ruget F., Andrianorisoa S., Le Bas C., Justes E., Léonard J. (2015). Accuracy, robustness and behavior of the STICS 8.2.2 soil-crop model for plant, water and nitrogen outputs: evaluation over a wide range of agro-environmental conditions in France. Environmental Modelling & Software, 64, 177-190 |
| SUCROS | Expert-N 3.0 | van Laar, H.H. , J. Goudriaan, und H. van Keulen, 1992: Simulation of crop growth for potential and water-limited production situations (as applied to spring wheat).: Simulation Report CABO-TT no. 27. Wageningen: Centre for Agrobiological Research and Department of Theoretical Production Ecology, Wageningen Agricultural University;<br><br>Vanclooster, M., Viaene P., Diels J., Christiaens K., 1994: WAVE a mathematical model for simulating water and agrochemicals in the soil and vadose environment. Reference and user's manual (release 2.0). Leuven: Institute for Land and Water Management, Katholieke Universiteit Leuven. |
| PCWOFOST | 5.3.3 | Ceglar A. , van der Wijngaart R. , de Wit A., Lecerf R., Boogaard H., Seguini L. , van den Berg M., Toreti A., Zampieri M., Fumagalli D., Baruth B. (2019). Improving WOFOST model to simulate winter wheat phenology in Europe: Evaluation and effects on yield. Agricultural Systems. 168, 168-180. |
| WCCWOFOST | 7.1.7 | Boogaard, H.L., Van Diepen, C.A., Rötter, R.P., Cabrera, J.M.C.A., Van Laar, H.H., 1998. User's guide for the WOFOST 7.1 crop growth simulation model and WOFOST control center 1.5. Technical Document 52. Winand Staring Centre, Wageningen, the Netherlands, 144 pp. |
| Wheat-Grow | 3.1 | Zhu Y.; Liu L.; Liu, B. WheatGrow: A simulation model for predicting growth and productivity in wheat. In Proceedings of the Workshop on Modeling Wheat Response to High Temperature, Texcoco, Mexico, 19–21 June 2013. |

| | | |
|---|---|---|
| | | Lv Z., Liu X., Tang L., Liu, L., Cao, W. and Zhu, Y., 2016. Estimation of ecotype-specific cultivar parameters in a wheat phenology model and uncertainty analysis. Agricultural and Forest Meteorology, 221: 219-229. |

**Table S1**

**Model structures used in this study**

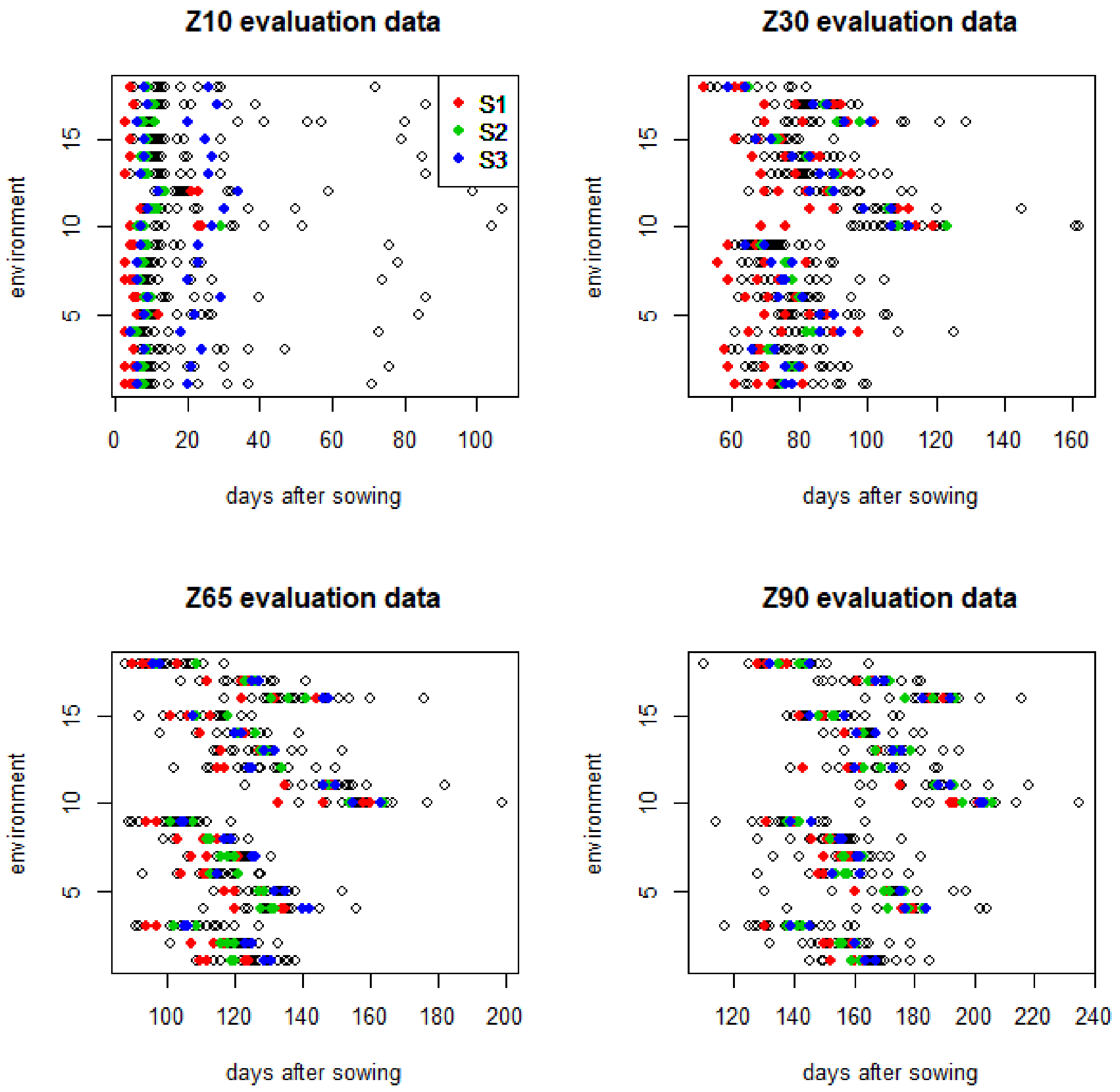


**Figure S1**

**Predictions of days from sowing to Zadoks stages Z10 (emergence), Z30, Z65 and Z90 by each modeling group for each evaluation environment. Modeling groups that used the same model structure are identified by color (red for structure S1, green for structure S2, blue for structure S3).**

| | MAE_eval | MAE_Z30 | MAE_Z65 | MAE_Z90 | RMSE_eval | EF_eval | skillT_eval | MAE_cal |
|---|---|---|---|---|---|---|---|---|
| M9 | 6.3 | NA | 9.3 | 3.2 | 7.2 | 0.7 | 0.489 | 6.2 |
| emean | 6.3 | 8.8 | 7.3 | 2.9 | 8.1 | 0.64 | 0.38 | 6 |
| M24 | 6.4 | 9 | 6.8 | 3.2 | 8.6 | 0.62 | 0.351 | 8.5 |
| emedian | 6.4 | 8.6 | 7.4 | 3.3 | 8.3 | 0.63 | 0.367 | 5.9 |
| M21 | 6.6 | 8.7 | 6.6 | 4.4 | 8.5 | 0.64 | 0.379 | 5.9 |
| M4 | 6.7 | 9.8 | 6.4 | 3.8 | 8.4 | 0.65 | 0.39 | 5.7 |
| M2 | 6.8 | 10.4 | 7.3 | 2.8 | 8.8 | 0.57 | 0.263 | 6.3 |
| M13 | 7.2 | 10.6 | 7.9 | 3.2 | 9 | 0.55 | 0.231 | 8.3 |
| M18 | 7.2 | NA | 10.8 | 3.6 | 8.6 | 0.62 | 0.336 | 7.7 |
| M15 | 7.3 | 11.7 | 4.6 | 5.6 | 8.5 | 0.64 | 0.383 | 8 |
| M11 | 7.3 | 7.7 | 7.4 | 6.8 | 10 | 0.53 | 0.18 | 9 |
| M25 | 7.4 | 10.7 | 6.7 | 4.8 | 8.9 | 0.6 | 0.307 | 6.1 |
| M23 | 7.8 | 10.6 | 8.5 | 4.2 | 10 | 0.5 | 0.14 | 6.7 |
| M26 | 7.9 | 9.3 | 10.3 | 4.2 | 10.3 | 0.47 | 0.082 | 7.4 |
| M3 | 8 | NA | 10.1 | 6 | 9 | 0.61 | 0.322 | 7.7 |
| M27 | 8 | 9.7 | 10.9 | 3.4 | 10.1 | 0.46 | 0.066 | 6.7 |
| M29 | 8.2 | 8.2 | 8.1 | NA | 11.2 | 0.44 | 0.029 | 9.7 |
| onlyT | 8.2 | 10.7 | 10.6 | 3.2 | 10.5 | 0.42 | 0 | 8 |
| M17 | 8.4 | 7.3 | 7.3 | 10.6 | 10.4 | 0.52 | 0.165 | 7 |
| M19 | 8.5 | 11.4 | 10.2 | 3.8 | 10.4 | 0.44 | 0.032 | 7.9 |
| M12 | 9.3 | 12.4 | 7.3 | 8 | 11.4 | 0.39 | -0.058 | 8.1 |
| M7 | 9.3 | 15.4 | 9 | 3.4 | 11.5 | 0.23 | -0.323 | 13.3 |
| M20 | 9.3 | NA | 11.5 | 7.2 | 11.4 | 0.4 | -0.041 | 8.3 |
| M8 | 9.4 | 12.4 | 8.8 | 6.8 | 11.6 | 0.35 | -0.117 | 12 |
| M22 | 9.5 | 9.1 | 15.8 | 3.4 | 12.1 | 0.21 | -0.362 | 7.3 |
| M10 | 10.6 | 24.8 | 5.9 | 1 | 13.3 | -0.54 | -1.664 | 12 |
| naive | 11.3 | 12.2 | 14.3 | 7.5 | 14.6 | 0 | -0.727 | 17.1 |

| | | | | | | | | |
|---|---|---|---|---|---|---|---|---|
| M14 | 13 | 15.2 | 13.3 | 10.5 | 16.2 | -0.18 | -1.044 | 18.7 |
| M6 | 14 | 9.9 | 10 | 22 | 17.6 | -0.47 | -1.537 | 8.1 |
| M16 | 14.2 | 11.8 | 10 | 20.8 | 16.6 | -0.29 | -1.228 | 13.9 |
| M1 | 15.8 | 20.6 | 10.4 | 16.4 | 16.8 | -0.33 | -1.301 | 12.8 |
| M28 | 20 | 18.3 | 24.2 | 17.4 | 23.5 | -1.51 | -3.343 | 17.4 |

**Table S2**

**Prediction errors for each modeling group and for the models e-mean, e-median, naive and onlyT. The columns are MAE averaged over the stages Z30, Z65 and Z90 for the evaluation environments (days), MAE for each of the stages Z30, Z65 and Z90 for the evaluation environments (days), root mean squared error (RMSE) averaged over the stages Z30, Z65 and Z90 for the evaluation environments (days), the skill measures EF and skillT averaged over the stages Z30, Z65 and Z90 for the evaluation environments (unitless) and MAE averaged over the stages Z30, Z65 and Z90 for the calibration environments (days). The models are ordered by average MAE (value in first column). NA indicates that that modeling group didn't predict the time to the indicated stage.**

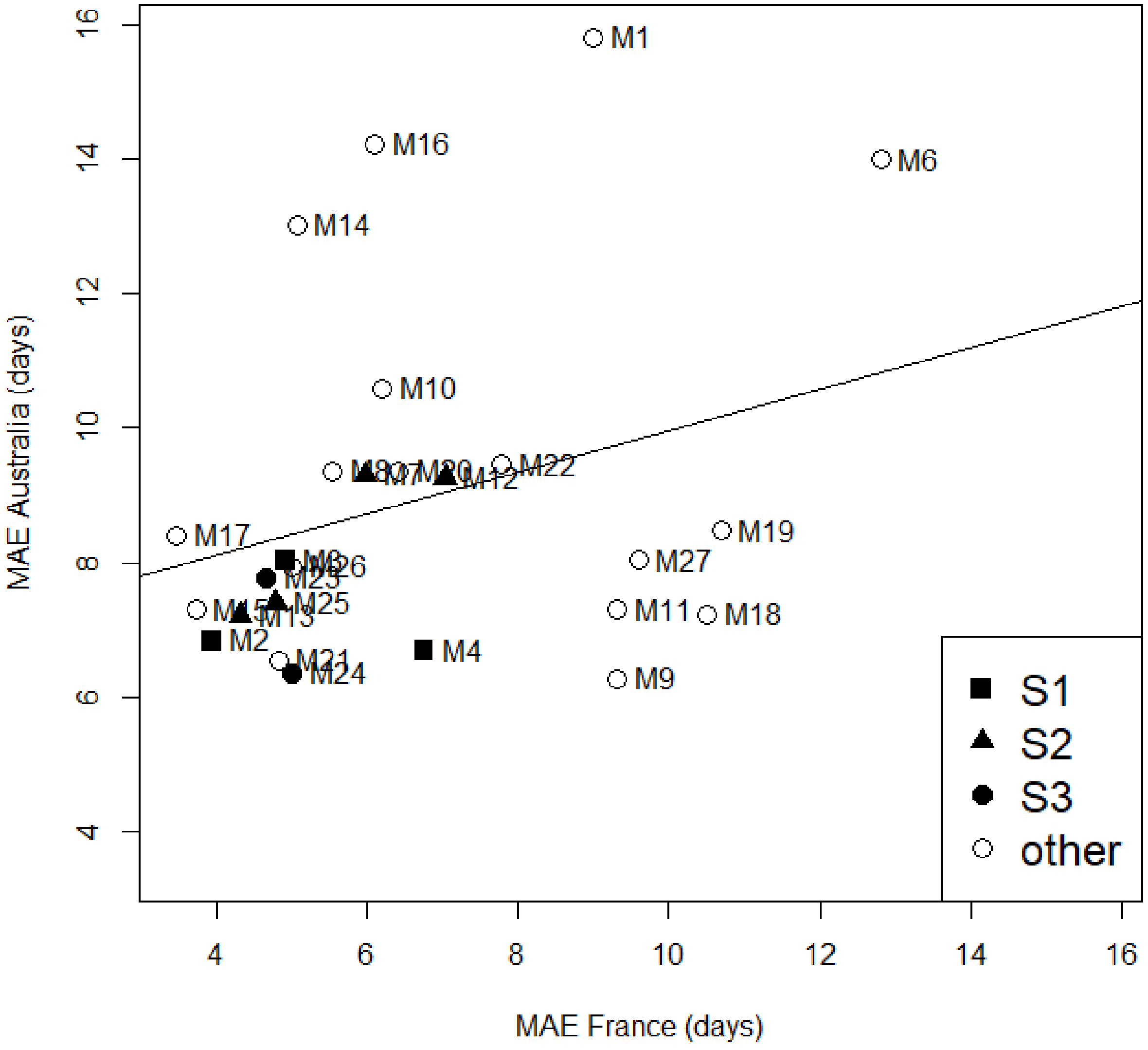


**Fig. S2**

**Relation between mean absolute error (MAE) for the Australian environments and MAE for the French environments, for modeling groups that participated in both studies. Values are averages over predicted development stages. Points are identified by modeling group. Modeling groups that shared the same structure (S1, S2 or S3) are identified by filled squares, triangles or circles, respectively. The regression line is y=8.23+0.24x.**
